# On metrics for subpopulation detection in single-cell and spatial omics data

**DOI:** 10.1101/2024.11.28.625845

**Authors:** Siyuan Luo, Pierre-Luc Germain, Ferdinand von Meyenn, Mark D. Robinson

## Abstract

Benchmarks are crucial to understanding the strengths and weaknesses of the growing number of tools for single-cell and spatial omics analysis. A key task is to distinguish subpopulations within complex tissues, where evaluation typically relies on *external* clustering validation metrics. Different metrics often lead to inconsistencies between rankings, highlighting the importance of understanding the behavior and biological implications of each metric. In this work, we provide a framework for systematically understanding and selecting validation metrics for single-cell data analysis, addressing tasks such as creating cell embeddings, constructing graphs, clustering, and spatial domain detection. Our discussion centers on the desirable properties of metrics, focusing on biological relevance and potential biases. Using this framework, we not only analyze existing metrics, but also develop novel ones. Delving into domain detection in spatial omics data, we develop new external metrics tailored to spatially-aware measurements. Additionally, a Bioconductor R package, poem, implements all the metrics discussed. While we focus on single-cell omics, much of the discussion is of broader relevance to other types of high-dimensional data.

## Introduction

A critical feature of single-cell omics data is its ability to elucidate heterogeneous cell subpopulations within complex tissues. Identifying and distinguishing these subpopulations is a fundamental task in single-cell analysis, often involving clustering cells into meaningful groups based on the similarity of their measurements. This process is typically preceded by several preprocessing steps, including normalization, feature selection, dimensional reduction, and batch correction, and the results of clustering serve as an entry point for assessing the performance of these upstream computational tasks [1–5]. Thus, evaluating the effectiveness of cell subpopulation identification is crucial for informing current methodological progress in single-cell data analysis.

When assessing the performance of cell subpopulation identification, researchers usually rely on external labels (the so-called ground truth) derived either from orthogonal experimental approaches (such as fluorescence-activated cell sorting, or additional data modalities in multiomics data) or from simulation. Validation metrics are then employed to quantify the degree of concordance between the clustering outcomes and these labels (also known as external clustering metrics). These metrics define the criteria for the quality of cell subpopulation identification, and therefore shapes the field’s methodological development.

However, conducting reliable and biologically meaningful performance evaluation remains challenging. Firstly, despite extensive discussions on external validation indices for general clustering tasks [6–8], there are no specific guidelines for choosing metrics in the context of single-cell biology. Researchers often follow established practices from the literature that, as we will show, may not always be optimal or well-justified. Secondly, while using multiple metrics for benchmarking is generally advisable, it can lead to inconsistent rankings of methods [3], making it difficult to draw clear conclusions. Finally, cellular heterogeneity in biology is often organized into complex hierarchical structures, but the ground-truth labels we use typically represent only one level of this hierarchy. As we will show, evaluations that expect a match strictly at this single level, without allowing for differences in resolution, can introduce bias that misrepresents the actual biological interest.

Inspired by previous research efforts to define formal constraints, or “desirable properties”, for external clustering metrics [6, 7, 9], we structure our discussion of various metrics around the key properties most relevant for single-cell biology. We seek to provide not only guidelines for selecting metrics and interpreting benchmarking results, but also a framework for developing guidelines applicable to new metrics and related computational tasks.

In the final section, we demonstrate how desirable properties and corresponding metrics can be adapted for spatial domain detection. Moreover, we show that the insights gained from clustering tasks can be applied as heuristics, and, when combined with spatial considerations, can inform the development of new desirable properties and metrics. Overall, it highlights how our framework can generate valuable guidelines not only for choosing appropriate metrics, but also for developing novel metrics that can better serve the research questions.

## Material & Methods

### A Datasets

#### A.1 The 10XPBMC dataset used in Figure 2 and Figure S3

This dataset is the scATAC-seq part of the 10XPBMC multiomics datasets used in [4], and the details about its generation, quality control (QC), and preprocessing steps can be found there. The clusterings in Figure 2 is done using the pipeline SnapATAC, and the clusterings in Figure S3 is done using the pipeline aggregation. See [4] for the pipeline details.

**Figure 1.**
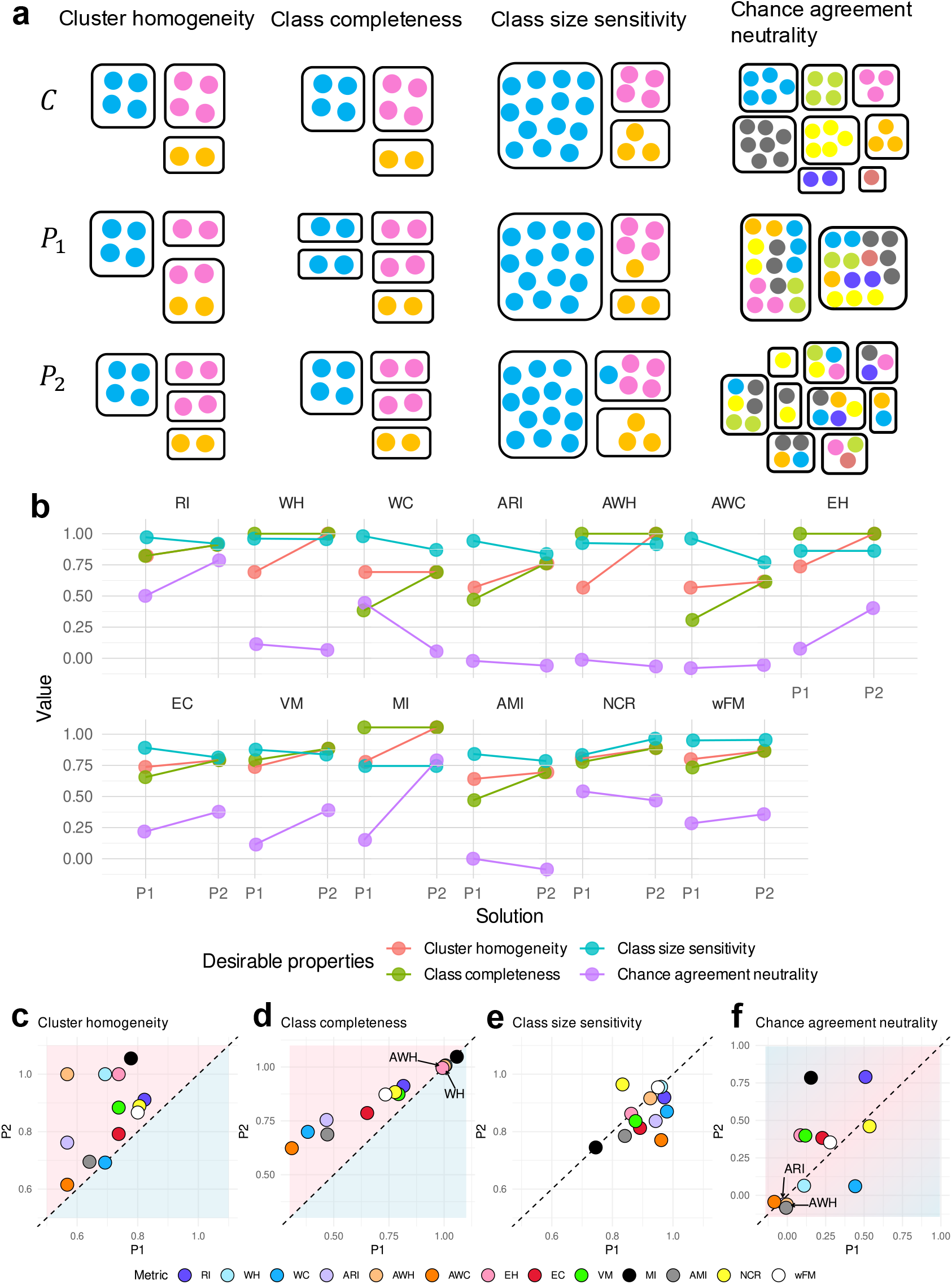
Desirable properties and how partition-based metrics reflect each property. Here in **a**, *C* represents ground truth partitions, while *P*_1_ and *P*_2_ are two hypothetical clustering results. In the first and second columns, *P*_2_ should be preferred to *P*_1_ according to the desirable properties of cluster homogeneity and class completeness, respectively. In the third column, if errors in smaller classes should be penalized more than errors in larger classes, then *P*_2_ should be preferred over *P*_1_, and vice versa. In the last column, *P*_1_ and *P*_2_ are two random partitions into different number of clusters. According to the property of chance agreement neutrality, neither *P*_1_ nor *P*_2_ should be preferred over the other. **b** illustrates the effectiveness of each metric in comparing *P*_1_ to *P*_2_ with respect to the corresponding properties. The values are calculated using the toy examples in **a. c**-**f** summarizes the comparisons from a property-centric perspective.

**Figure 2.**
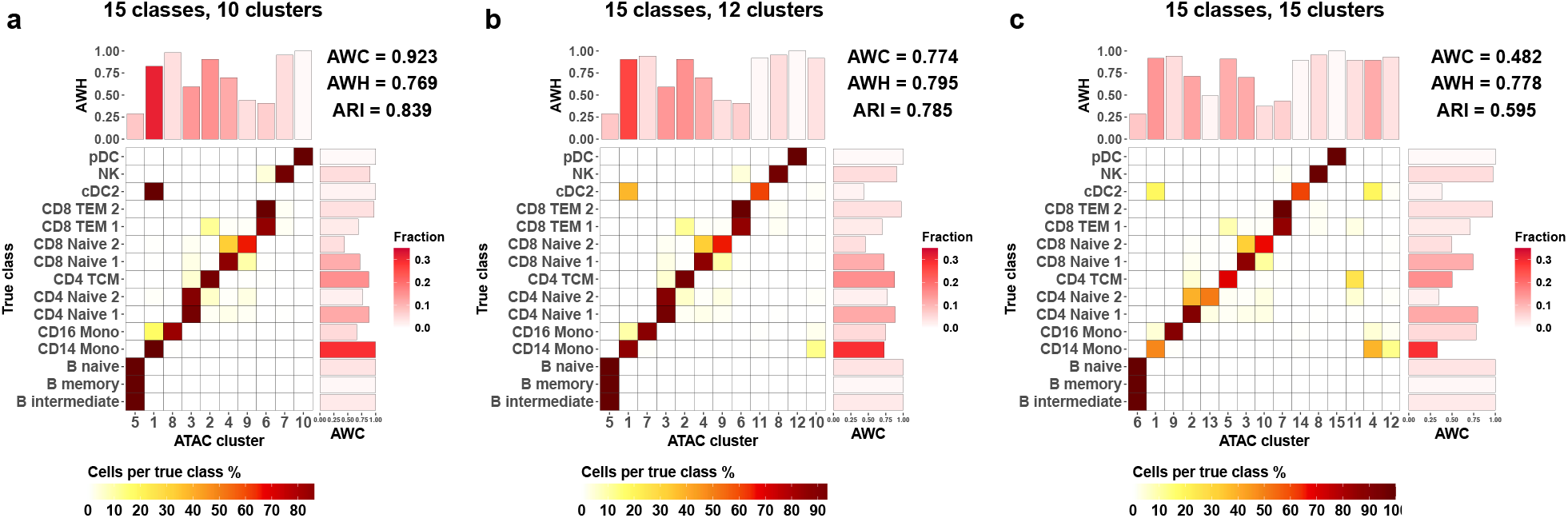
Real-world examples of partitions with different rankings across metrics. As an example, we use the 10X multi-omic PBMC dataset in Luo et al. [4], comparing three ATAC-seq clustering solutions against RNA-seq annotations (treated as ground truth). The x-axis represents the predicted clusters; y-axis is the ground truth classes. The tile colors indicate the proportion of cells from the corresponding true class (each row sums up to one). A clearer diagonal structure indicates better agreement. AWH, AWC and ARI values are shown. The barplot on top shows cluster-wise AWHs (homogeneity); the barplot on the right shows class-specific AWCs (completeness). Bar color represents the proportion of cells in each cluster/class.

#### A.2 The Atlas2 dataset used in Figure 4

See B.

**Figure 3.**
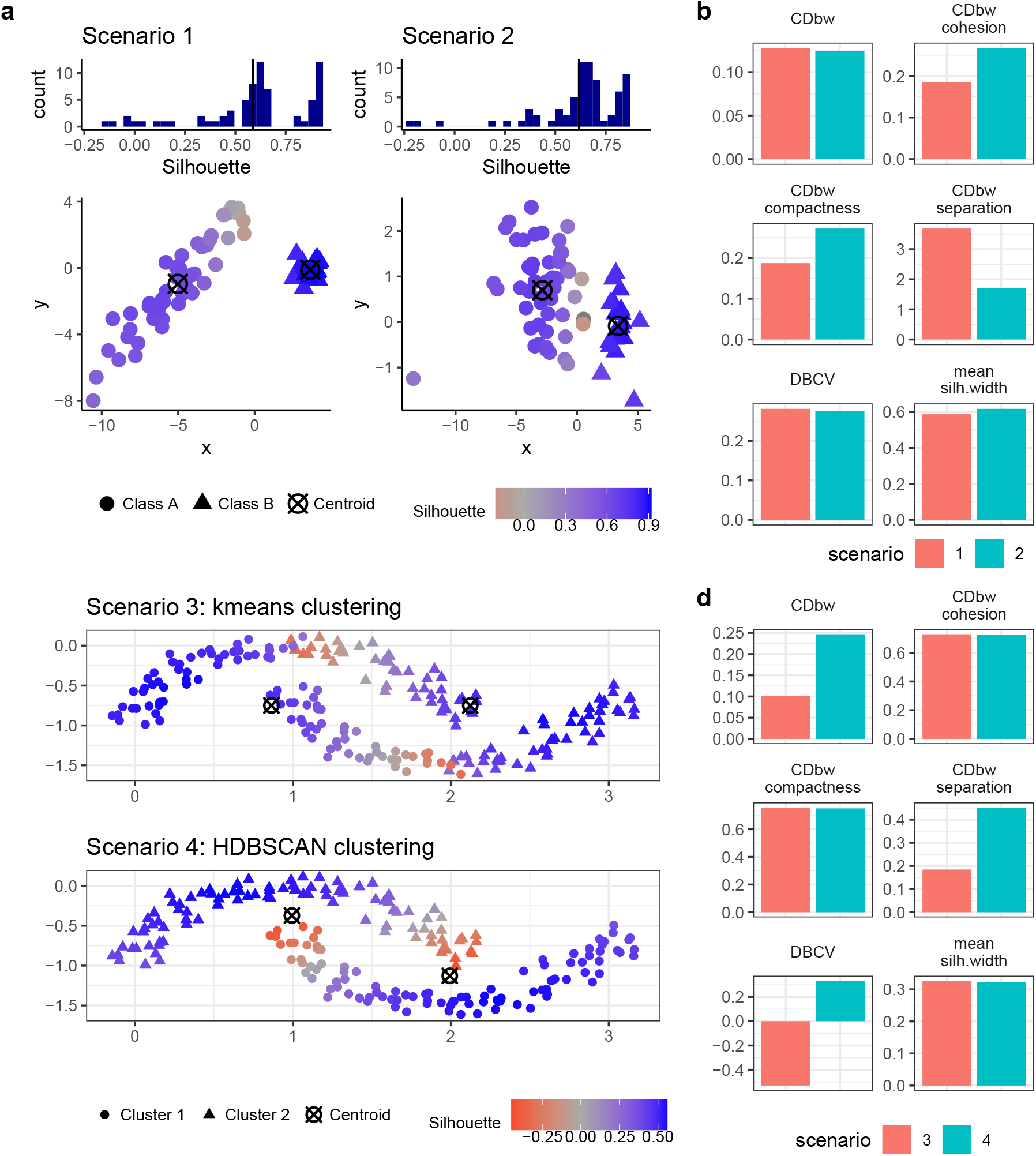
Examples of how class shape affects the embedding-based metrics. In **a**, both scenarios contain two classes sampled from 2D Gaussian distributions. Class A in Scenario 1 forms an elongated shape, while in Scenario 2, it is more globular. Silhouette scores for these two scenarios reveal similar distributions, despite the visually poorer separation of the two classes in Scenario 2. **b** illustrates how embedding-based metrics compare Scenario 1 to Scenario 2. In **c**, Scenario 3 and 4 highlight a moon-shaped dataset colored by clustering predictions from kmeans clustering and HDBSCAN clustering, respectively. **d** illustrates how embedding-based metrics compare Scenarios 3 and 4.

**Figure 4.**
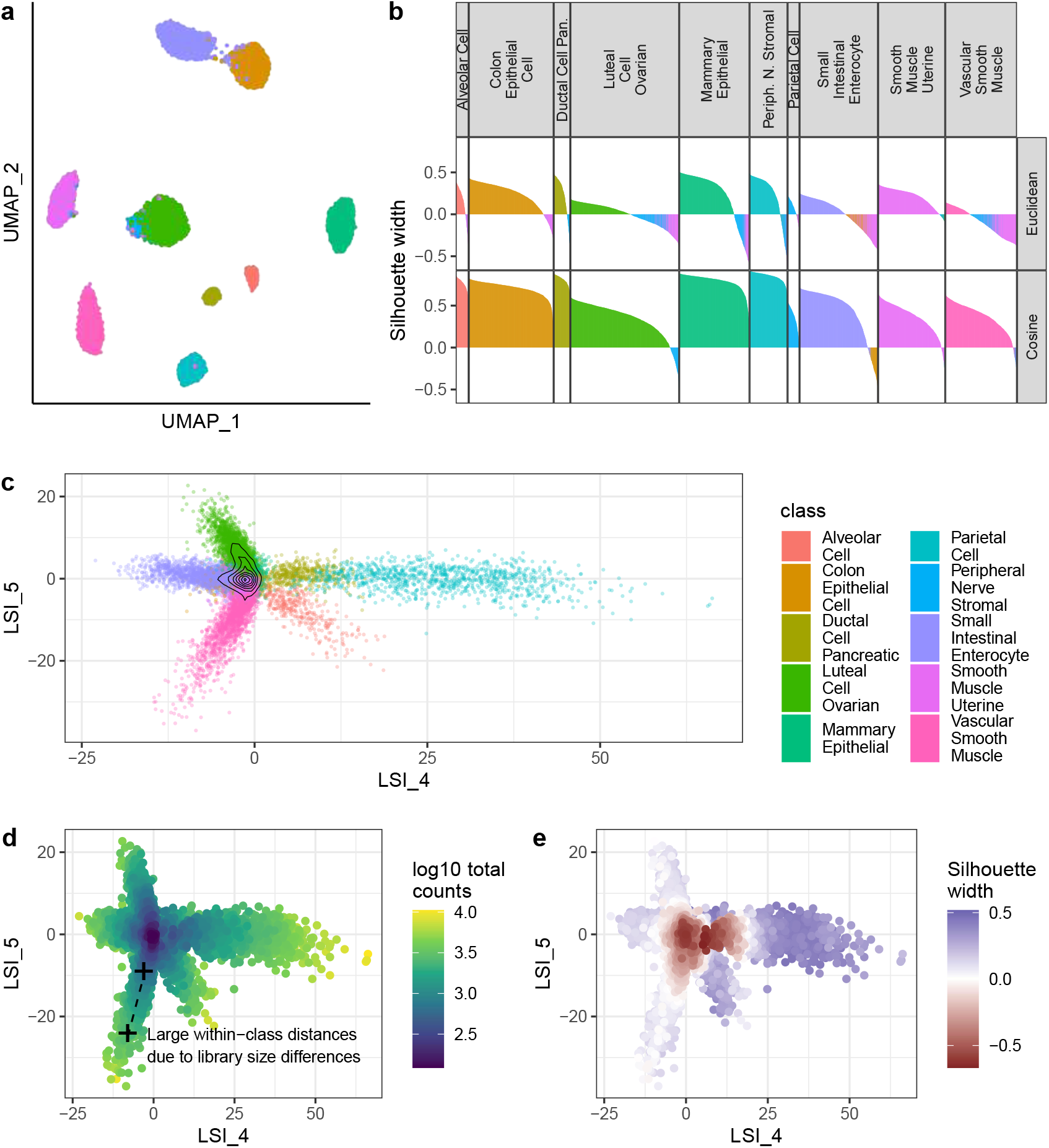
Real-world example of misleading Silhouette scores and Euclidean distances. Using the Atlas2 dataset from Luo et al. [4], UMAP embeddings (**a**) show well-separated classes. However, Silhouette scores using Euclidean distance in the LSI space (upper panel of **b**) show many cells with low scores; using Cosine distance strongly mitigates this effect (lower panel of **b**). Cells in **b** are colored with the identity of its own class if its silhouette width is positive and that of the closest other class if the width is negative. The LSI embedding in **c**, focusing on two LSI dimensions that best distinguish vascular smooth muscle cells—those with the highest proportion of negative Silhouettes in **a**—exhibits a star-shaped distribution (**c**). **d** shows library size for each cell, where higher values increase divergence between cell types and amplify within-class distances, collectively contributing to negative Silhouette scores (**e**).

#### A.3 The 10X Visium dataset

We use the LIBD 10X Visium dataset (slice 151673) from Maynard et al. [10], with manual annotations from the original publication serving as the ground truth. Domain detection is performed using BayesSpace, CellCharter, GraphST, precast, and SEDR with default parameters and specifying the true number of classes. Then the Hungarian algorithm is applied to match the predicted clusters with the annotated classes.

### B Exploration of the LSI embeddings for the Atlas2 scATAC-seq dataset

This is related to Figure 4. The dataset is the scATAC-seq Atlas2 dataset used in [4], and the details about its generation, quality control (QC), and preprocessing steps can be found there. After QC and filtering, the pipeline ArchR_tiles from [4] is used for dimensional reduction, which generates the latent sementic indexing-based (LSI) embeddings for each cell. Following ArchR’s documentation, the first LSI component is removed due to its above 0.75 Pearson’s correlation with the library size. The UMAP in Figure 4a and the Silhouette width in Figure 4b are calculated based on the 2^nd^ to 16^th^ LSI components.

### C Composed Density between and within Clusters (CDbw)

Let *C*_*K*_ denotes a clustering to be evaluated. Firstly, a number of representative points are selected for each cluster in *C*_*K*_ using a furthest-first technique to cover the geometry of the cluster effectively. The CDbw index is defined as

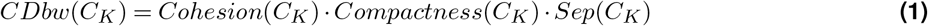

where

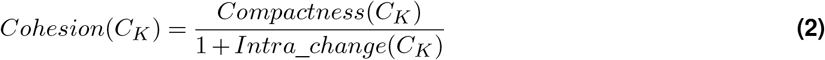

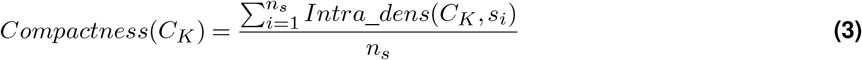

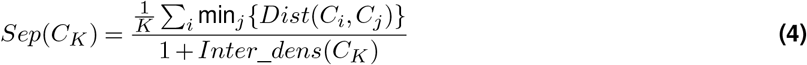

*Intra*_*dens*(*C*_*K*_, *s*_*i*_) is the intra-cluster density, and *s*_*i*_, *i* ∈ {1, …, *n*_*s*_} is a sequence of shrinking factors [0, 1] that is used to move the representative points closer to the cluster center. For a given *s*_*i*_, *Intra*_*dens*(*C*_*K*_, *s*_*i*_) calculates the average density around all the representative points, normalized by the standard deviation of the cluster. *Intra*_*change*(*C*_*K*_) is the changes in the within-cluster densities over changes in *s*_*i*_, which averages the difference between values of *Intra*_*dens*(*C*_*K*_, *s*_*i*_) for adjacent values of *s*_*i*_. To calculate *Sep*(*C*_*K*_), each representative of a cluster is paired with the closest representatives of any other cluster. *Dist*(*C*_*i*_, *C*_*j*_), the distance between clusters *C*_*i*_, *C*_*j*_, is calculated as the average of distances between the pairwise closest representatives; *Dens*(*C*_*i*_, *C*_*j*_), the density between clusters *C*_*i*_ and *C*_*j*_, is an average standardized number of points in the neighborhood of the midpoint of the line between each pair of representatives; The density between clusters in the clustering, *Inter*_*dens*(*C*_*K*_), is the average of maximum *Dens*(*C*_*i*_, *C*_*j*_) for every pair of clusters *C*_*i*_ and *C*_*j*_.

### D Density Based Clustering Validation index (DBCV)

For a clustering solution *C*, the DBCV index is the weighted average of the validity indices of all clusters:

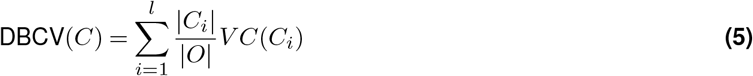

where *l* is the number of clusters, and |*O*| is the total number of objects. The validity index for a cluster *C*_*i*_ is calculated as the following:

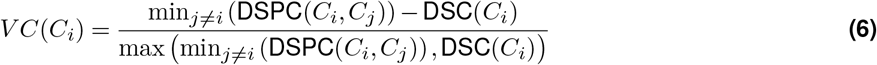

DSPC(*C*_*i*_, *C*_*j*_) is the density separation between two clusters *C*_*i*_ and *C*_*j*_, which is defined as the minimum mutual reachability distance between their internal nodes:

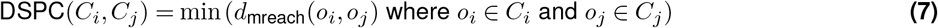

whereas DSC(*C*_*i*_) is the density sparseness of a cluster *C*_*i*_, being the maximum edge weight (largest distance) of the internal edges in the Minimum Spanning Tree (MST) of the cluster. The MST is constructed using the mutual reachability distance considering the objects in *C*_*i*_.

*d*_mreach_(*o*_*i*_, *o*_*j*_) is the mutual reachability distance for two objects *o*_*i*_ and *o*_*j*_, and is defined as:

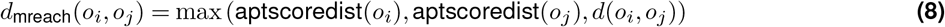

where *d*(*o*_*i*_, *o*_*j*_) is the pairwise distance between objects *o*_*i*_ and *o*_*j*_, and aptscoredist(*o*_*i*_) is the core distance of object *o*_*i*_. For an object *o* in a cluster *C*_*i*_, its core distance with respect to all other objects in the same cluster is calculated by

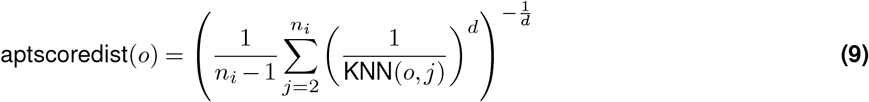

where: *n*_*i*_ = |*C*_*i*_|, KNN(*o, j*) is the distance to the *j*-th nearest neighbor of object *o* within the same cluster, and *d* is the dimensionality of the feature space.

### E Entropy-based Local indicator of Spatial Association (ELSA)

For a site *i*, the ELSA score is the product of two terms:

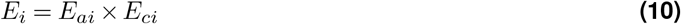

where *E*_*ai*_ summarizes the dissimilarity between site *i* and the neighbouring sites and is calculated as

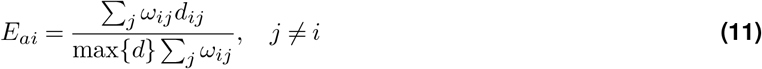

*ω*_*ij*_ is a binary weight that specifies whether the site *j* is within the neighbourhood of site *i*, and *d*_*ij*_ describes the dissimilarity between the categories at sites *i* and *j*, which is calculated as the absolute difference of the ranks assigned to the categories at sites *i* and *j*.

*E*_*ci*_ on the other hand quantifies diversity of the categories within the neighbourhood of site *i*:

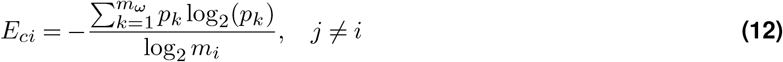

where *m* are the total number of categories, *p*_*k*_ is the probability of *k*th category from the *m*_*ω*_ categories within the neighborhood of site *i*, and 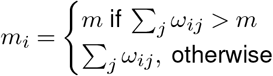.

### F Simulations

This is related to Figure S5. To explore in a mostly unbiased way the behavior of metrics, we generate a number of simulated 2- and 3-dimensional datasets with 3 to 4 classes of varying abundances (some equal, some highly skewed), variability, and differences between classes. This is implemented in the `mockData` function of the `poem` package. Briefly, we first use multi-dimensional scaling to obtain centroids for each class that approximate the desired pairwise class differences (determined by the settings), and then generate the desired number of points around them using a normal or log-normal distribution with a given variability parameter. To create further non-gaussian spread in some classes, we concatenate points from multiple such distributions symmetrically shifted from the centroid. Each simulation was further performed with 2 seeds (see the github repository for the exact sets of parameters used), yielding 2240 simulations. For each, we then take the simulated data to be embedding-like strucutres, and calculate embedding-level metrics as well as graph metrics on a kNN graph with k=5. To compute partition-level metrics, we run k-means clustering with k ranging from 2 to 6, as well as Louvain clustering on the kNN graph with resolutions of 0.5, 1 and 2. Node- and class-specific metrics are averaged to yield global metrics. To enable a comparison of the metrics results with interpretable dimensions of the simulation parameters, we define “size imbalance” as the Simpson diversity index of the class abundances, the “signal-to-noise” as the ratio between the mean pairwise class differences and the mean class variability (i.e. standard deviation), and the “spread” as the mean class spread parameter. We then perform a linear regression of each metric on these three parameters across simulations, and for partition-level metrics we additionally include the number of clusters as a covariate.

### G Fuzzy pair-counting metrics

We use Hullermeier’s Normalized Degree of Concordance (NDC, see Hullermeier et al. [11]) as a fuzzy version of the RI, and d’Ambrosio’s Adjusted Concordance Index (ACI, see D’Ambrosio et al. [12]) as fuzzy ARI. The ACI adjusts the NDC for chance based on permutation of the membership probability vectors. Based on this logic, we further implemented fuzzy Wallace indices (i.e. Wallace Homogeneity and Wallace Completeness and their chance-adjusted versions). Briefly, Wallace Completeness of a class is originally defined in the non-fuzzy setting as the proportion of all the pairs of elements of that class that are also in the same cluster. In the fuzzy setting, this pair concordance is understood as the similarity in membership agreements for pairs of elements between the two fuzzy partitionings. Specifically, the agreement between a pair of elements *i* and *j* is given by 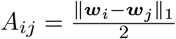, where ***w***_*i*_ and ***w***_*j*_ represent their respective membership vectors. The concordance between two fuzzy partitionings *Q* and *P* for that pair is then given as 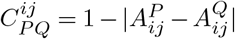. Since class membership is fuzzy, the extent to which this concordance matters for the computation of the completeness of a class *X* depends on the extent to which the pair of elements are of that class, which can be computed using different *t*-norm operations for set membership. While multiple such functions are implemented in our package, by default we use the product *t*-norm, meaning that the degree of membership of a pair of element to the class *X* (denoted 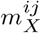) is given by the product of the probability that each of the two element belongs to class *X*. The Wallace Completeness of class *X, WC*_*X*_, can then be computed as 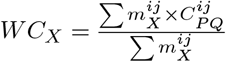 across all possible pairs *i* and *j*. To obtain a chance adjusted *AWC*_*X*_, we compute the randomly expected 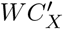 across permutations of the predicted memberships, and then calculate 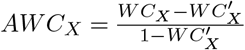. The same is done for Wallace Homogeneity.

### H Spatial application of fuzzy metrics (Neighborhood-smoothed pair sorting metrics)

To make hard labels fuzzy in a spatially-dependent fashion, we first compute the domain composition of its spatial nearest neighbors (i.e. nearest neighbors based on the Euclidean distance between spots/cells). To make the neighborhood definition more comparable across areas, in particular at the border of a field of view (where there are fewer first-degree neighbors) or in areas of low-density for non-spot-based datasets, we compute the median distance to the desired *k*^*th*^ nearest neighbor across spots/cells, and then consider for each spot/cell the neighbors that are within that range. The neighborhood domain membership (***w***_n_) of a spot *i*, ***w***^*i*^, is then the number of its neighbors belonging to domain, divided by the number of neighbors. The domain membership of spot *i* itself, 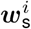, is 1 if the spot belongs to a domain, and zero otherwise. The fuzzy domain membership is then computed as 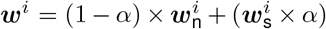, where *α* is a parameter controling the relative weight of the spot versus it’s neighborhood (by default, we set *α* = 0.5).

By comparing against a fuzzy truth, we decrease the concordance of a predicted partitioning that would be exactly like the original ‘hard’ truth. For this reason, we developed a fuzzy-hard version of the metrics that compares (hard) clustering labels to both a “hard” and a fuzzy ground truth. For the purpose of computing within-pair agreements, the hard labels are expressed as a weight of 1 for the given label and 0 for other labels. Otherwise the procedure is exactly like described above, except that the concordance between the prediction *P* and the truth *Q* of a pair of elements *i* and *j* is given by 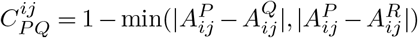, where *Q* and *R* are respectively the hard and fuzzy versions of the ground truth. In this context, for the purpose of computing completeness, pair membership is based on the hard ground truth labels.

### I Spatial RI/ARI

Yan et al. [13] proposed a spatially-aware version of the Rand Index (RI) and its adjusted ARI. In this context, corcordant pairs (i.e. pairs that are either separated or grouped according to both partitions) retain a concordance score of 1, as in the classical RI. However, discordant pairs, which would score 0 in the classical RI, get assigned a certain value depending on the distance between the two spots/elements. How this value is set depends on the type of discordance. Specifically, if the elements are *wrongly grouped in the clustering* (i.e. but are separated in the ground truth), then this mistake is more tolerated if the elements are spatially close to each other, and the value is determined by the function *h*(*d*):

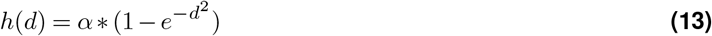

Instead, if the elements are *wrongly separated in the clustering* (i.e. but are grouped in the ground truth), then this mistake is more tolerated if the elements are spatially distant, and the value is determined by the function *f* (*d*):

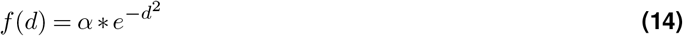

The distance, here, is computed on normalized (0-1) coordinates, and the authors set the *α* to 0.8.

As shown in Figure S10d-f, we find these original decay functions to be insufficiently steep: wrongly grouped elements get tolerated even if they are quite distant, and vice versa. We therefore changed the decay functions to the following:

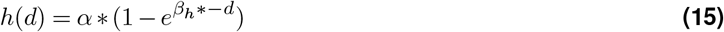

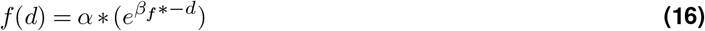

with the default values of *β*_*f*_ = 4 and *β*_*h*_ = 1. The modified decay can be visualized in Figure S10g-i. These parameters are modifiable arguments to the function, and users can decide to revert back to the original distance functions from Yan et al. [13].

### Scalability Assessment

We used the Visium HD colon cancer dataset from Oliveira et al. [14] for benchmarking (see Figure S11 for dataset details). Metric calculations were performed using both the manual annotations provided in the original study and the predicted domain partitions or cell embeddings from CellCharter [15].

CPU time (cpu_time) and peak memory usage (max_rss, maximum resident set size) of metrics was assessed by using the Snakemake’s built-in benchmarking function. The benchmarking pipeline was executed on a computing cluster, with 8 CPU cores and 100 GB of memory allocated per core (i.e., sbatch -c 8 `–mem-per-cpu=102400`), allowing up to 800 GB of total memory across the workflow.

## Results

### Understanding partition-based metrics to evaluate clustering

To distinguish from measurements at the upstream data representation level, we use the term “partition-based metrics” for external clustering metrics (See Glossary box). For clarity, we refer to the predicted partition to be evaluated as “clusters”, and the ground-truth partition as “classes”. The principle for defining desirable properties of clustering metrics can be summarized as follows: Given any pair of partitions of the same set, denoted as (*P*_1_, *P*_2_), where *P*_2_ is assumed to be a better clustering option than *P*_1_ according to some intuition, a metric *F* should adhere to the constraint that *F* (*P*_1_) *< F* (*P*_2_) (assuming that a higher value of *F* indicates a better clustering) [7]. Some of these constraints and intuitions are about features of the partition that should be reflected by the metric, while others can be about biases that should *not* influence the metric.

Rather than reviewing all desirable properties in the literature [6, 7, 9], we focus on properties that are most relevant to clustering molecular profiles of cells: *cluster homogeneity, class completeness, class size sensitivity*, and *chance agreement neutrality*. Cluster homogeneity states that clusters should not mix objects from different classes. Class completeness requires that all elements of a single class should not be distributed across different clusters. Class size sensitivity refers to how the misassignment of an element is evaluated relative to the size of the class it belongs to. We note that whether and what exact sensitivity to class size is desired depends on the context and research question. For instance, a metric might be considered desirable if it: i) is insensitive to class size [16], ii) penalizes errors more in larger classes than in smaller ones, or iii) penalizes errors more in smaller classes than in larger ones [6]. Lastly, chance agreement neutrality means that similarities between two partitions occuring purely by chance should not be given undue credit to inflate the final similarity score (see Glossary box or Albatineh and Niewiadomska-Bugaj [17]).

Partition-based metrics can be categorized into three groups based on their calculation: pair-counting, information theoretic and set matching measures. Extensive literature covers what they are and how they compare to each other [6–8, 16, 18–20]. Instead of reviewing them all here, we focus on key aspects from each category, as listed in Table 1, and examine how they capture the desirable properties (Figure 1). Such an examination can be easily extended to additional metrics.

**Table 1.**
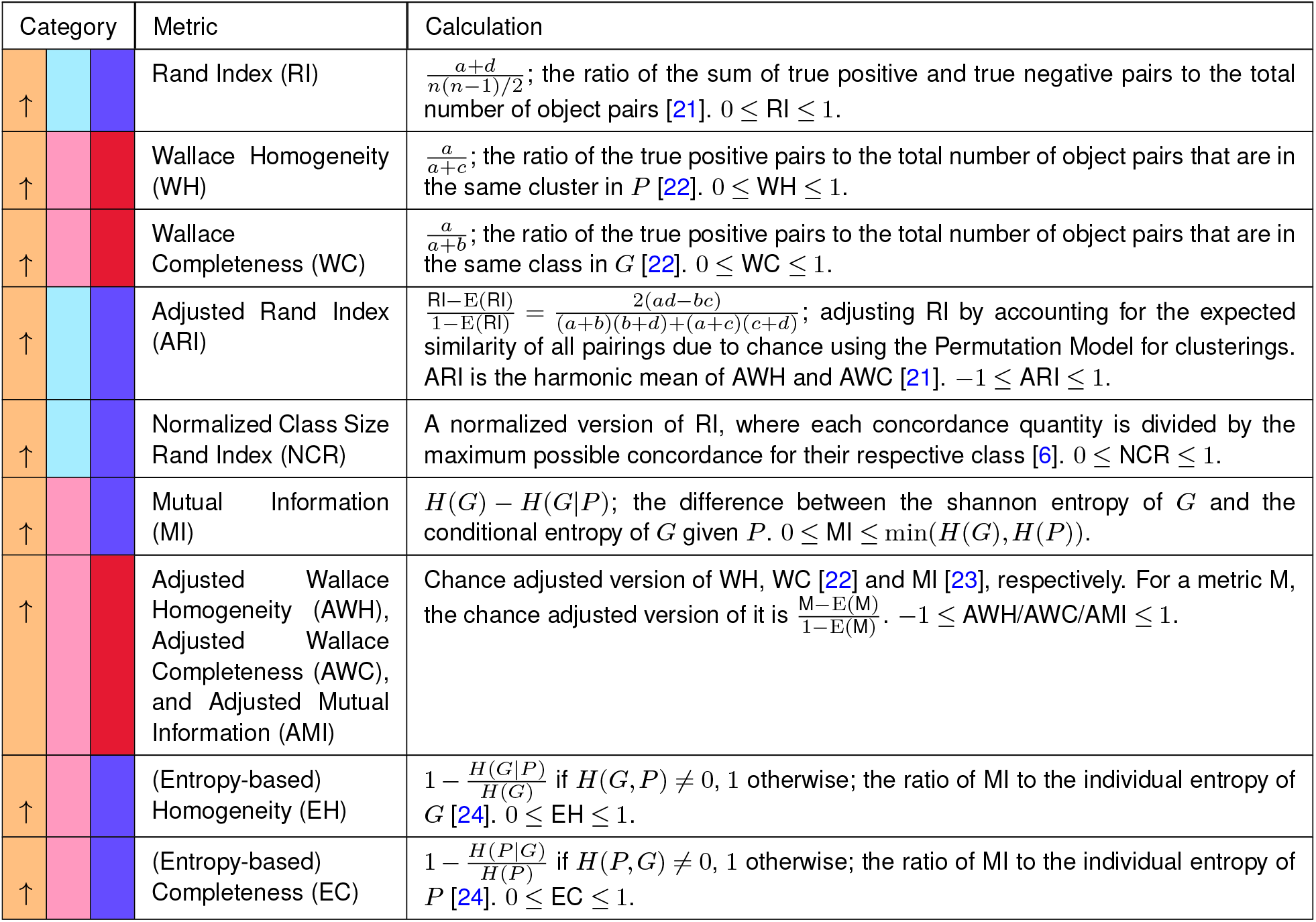
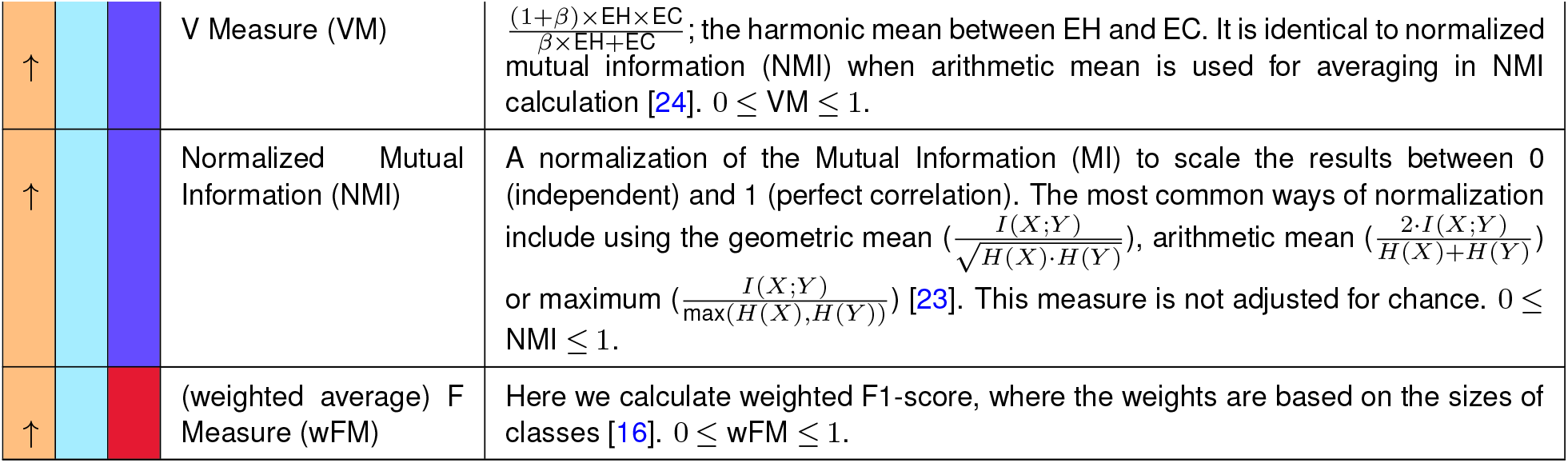
Partition-based metrics. Metrics are categorized according to whether they are aggregated or track a specific desirable property, and the minimal unit of calculation (cell, class/cluster, or dataset). In the first column, a ↑ indicates that a higher metric value is better, while a ↓ means that a lower value is prefered. The notation used is common throughout the table: consider comparing the predicted partition *P* to the ground-truth partition *G*; *a* is the number of pairs that are in the same group both in *P* and *G*; *b* is the number of pairs that are in the same class in *G* but in different clusters in *P* ; *c* is the number of pairs that are in different classes in *G* but in the same cluster in *P* ; *d* is the number of pairs that are in different groups both in *P* and *G*; *n* is the total number of objects; E is the expectation operator; *H*(*·*) is the Shannon entropy; *I*(*X*; *Y*) is the Mutual Information between *X* and *Y* ; *β* is the ratio of weight attributed to homogeneity vs completeness; the expectation values of RI, WH, and WC are calculated under a generalized hypergeometric model. 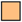 partition-based metric 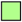 embedding-based metric 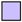 graph-based metric 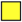 partition-based metric with spatial information 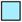 aggregated metric evaluation 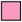 property-based metric 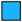 element-level evaluation 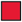 class/cluster-level evaluation 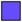 dataset-level

Based on how a metric reflects the properties of homogeneity and completeness, we categorize them into *property-based* metrics and *aggregated* metrics. Property-based metrics focus on a specific desirable property. Metrics that measure homogeneity include Wallace Homogeneity (WH), its adjusted version (AWH) [22], Mutual Information (MI), and Entropy-based Homogeneity (EH) [24]. Conversely, completeness metrics contain Wallace Completeness (WC), its adjusted version (AWC), and Entropy-based Completeness (EC). Aggregated metrics integrate both homogeneity and completeness. The Rand Index (RI) and its adjusted forms (Adjusted Rand Index, ARI, and Normalized Class Size Rand Index, NCR) are classic examples, with RI representing the harmonic mean of WH and WC, ARI being the harmonic mean of AWH and AWC, and NCR [6] a variant of ARI using a different normalization method (Table 1). Other key aggregated metrics include the V-measure (VM), which is the harmonic mean of EH and EC, and the (weighted average) F-measure (wFM) [16], which is the harmonic mean of precision and recall (the set-matching equivalents of homogeneity and completeness).

The use of toy examples in Figure 1a demonstrates how *P*_1_ or *P*_2_ should be preferred over the other according to each of the desirable properties. Figure 1b-f shows the metric values for these examples, illustrating their behavior. As expected, RI, ARI, NCR, wFM and VM effectively capture both homogeneity and completeness (Figure 1b), while WH, AWH, MI, EH reflect homogeneity independently of completeness changes. However, when MI is adjusted for chance (*i*.*e*. AMI), it can no longer be independent of completeness. WC, AWC and EC primarily reflect completeness, with AWC also showing a slight preference for *P*_2_ in the homogeneity example due to its chance-adjusted nature (See Glossary box), which accounts for the difficulty of achieving the same completeness with an increased number of clusters as in *P*_2_. Similarly, EC does not assign identical values to *P*_1_ and *P*_2_ in the homogeneity scenario because it lacks chance adjustment, making it sensitive to the number of clusters, particularly when both the number of samples and clusters are small [24].

Regarding class size sensitivity, Figure 1b and 1e illustrate that WH, AWH, MI, EH and wFM are insensitive to class size. NCR [6] is unique in its tendency to penalize errors in smaller classes more heavily. The remaining metrics tend to prefer errors in smaller classes over larger ones.

When it comes to chance agreement neutrality, metrics adjusted for chance, such as ARI, NCR, AWH, AWC and AMI (Figure 1b,f), effectively neutralize the influence of random agreements. In contrast, their unadjusted counterparts—RI, WC and MI—are biased, with the exception of WH. wFM seems to have some resilience to such bias. Other unadjusted metrics including EH, EC and VM, fail to remain neutral to random agreements. Interestingly, although both EC and WC measure completeness, they differ in their preference for *P*_1_ and *P*_2_, indicating that they can be affected differently by random agreements.

Notably, there is an inherent contradiction between the chance adjustment in metrics and a class size sensitivity that would favor the same error rate in larger classes over smaller ones. As depicted in Figure 1e, ARI gives greater advantage to *P*_1_ than *P*_2_ compared to RI, and similarly, AWC favors *P*_1_ over *P*_2_ more than WC does. Recall that both ARI and AWC are explicitly adjusted for chance. Such chance adjustment assumes that correct assignments in rare classes are more likely to happen by chance, so the adjusted score is reduced more for these cases (*e*.*g. P*_2_). However, this assumption may not always be valid in single-cell analysis, where methods that focus on major components of variability (e.g. SVD, variable feature selection, etc.) naturally make it challenging to detect rare populations. While the uniform sampling assumption is often acceptable for simplicity, in studies where rare populations are highly relevant, the difficulty of characterizing smaller classes should not be ignored. Thus, although chance adjustment is generally recommended for meaningful comparisons, it can introduce bias and might not necessarily track the most desired partitioning. When the accurate identification of rare classes is crucial, metrics such as NCR should be considered.

Overall, we show here how different metrics vary in their capability to reflect various desirable properties. This sheds light on why, in many benchmark studies, rankings often vary depending on the metric (Figure 2, [3, 4]). The present discussion and example analyses can therefore help users understand what each metric measures, guide their usage, and lead to better benchmark interpretation.

Furthermore, we want to highlight the limitations and lack of transparency associated with using aggregated metrics including ARI [25] and VM as a universal standard for assessing clustering performance. Although ARI can provide a general sense of clustering effectiveness, it does not always correlate with the optimal clustering strategy, particularly in complex tasks. We observed that as clustering granularity increases, a noticeable trade-off emerges between homogeneity and completeness, with higher granularity favoring homogeneity at the expense of completeness (Figure S1). This phenomenon is critical in contexts such as disease-relevant cell state discovery (Figure 2) or meta-cell identification (See Glossary box). While VM provides some flexibility by allowing adjustments to the weights between EH and EC, it remains unclear how to optimally tune this weight to achieve an optimal solution.

Therefore, despite the widespread use of aggregated metrics such as ARI, we advocate for a more selective reliance on them. While such metrics are useful for broadly identifying low-performing methods, more fine-grained comparisons among top-performing methods should instead focus on property-based metrics such as MI, AWH, and AWC, as they offer clearer insights into specific aspects of clustering performance. These metrics provide an easily interpretable readout of strengths and weaknesses, allowing for a more targeted approach in method selection and evaluation.

#### Challenges with partition-level evaluation

Evaluating at the partition level is straightforward, but determining a partition requires selecting the number of clusters, *k*, or parameters that influence it, and the values of partition-based metrics vary as *k* varies ([16, 18]; see also Figure S1). Multiple strategies have been adopted to address this problem, including evaluating the metric at the “best-performing *k*” [4], the “true *k*” (based on the available ground truth labels) [1, 2, 26], the “detected *k*” (typically using default parameters) [2], or across a range of parameters that yield different *k* values [4]. Each of these approaches, however, can introduce potential biases. For example, Figure 2a-c show clusterings generated by the same pipeline with different *k*. When using the “true *k*” (Figure 2c), the pipeline performance is underestimated, because certain populations, such as B cell subtypes, are very similar and difficult to distinguish. Forcing the clustering to match the “true *k*” results in unnecessary segregation of correctly identified clusters (Figure 2c), which significantly reduces AWC (completeness) without improving AWH (homogeneity). Thus, the clusterings based on the “true *k*” do not always represent the most relevant solution a pipeline can generate. Selecting the “best-performing *k*”, on the other hand, has the drawback that different metrics might not agree on the best *k*. This highlights the need for complementary evaluation approaches.

In addition, biological populations often exhibit a hierarchical structure [27], while the ground truth is typically available at a single resolution, which is not necessarily the only meaningful one. For example, within a population labeled as a single class by the ground truth, there may exist real and meaningful subpopulations. A method that identifies these subpopulations would be significantly penalized, because the partition itself does not carry any information regarding the relationships between the different classes or clusters. This issue has been discussed in Wu and Wu [27], where the authors proposed evaluation metrics for comparing a partition to a hierarchical ground-truth structure. As we will see, these issues can also be mitigated by evaluating at other levels of the analysis.

### Evaluations on upstream data representations

To circumvent the shortcomings of partition-level evaluations or at least to complement it with other measures, a promising strategy is to evaluate *upstream* aspects of the data, such as low dimensional embeddings or graph representations. This removes the need to select a specific *k* or resolution to explicitly define clusters. Moreover, because these representations can preserve relationships between clusters across resolutions (i.e. two subpopulations of a class will nevertheless tend to be close to each other), evaluating graph or embedding structures helps to mitigate biases related to parameter choices and the resolution of the ground truth. As we will see and as shown in the literature [28], however, it is important that the assumptions underlying the representations and metrics be aligned to the relevant downstream tasks.

Below, we first extend the desirable properties of partitions to embeddings and graphs. While formulations of these properties can be found in the literature [29, 30], they can be understood as adaptations of the general intuitions of homogeneity and completeness. At the embedding level, these include *compactness* and *separation* of the classes; at the graph level, they are *neighborhood homogeneity* and *class compactness*. The basic intuition is that elements of a given class should be close to each other (*i*.*e*. compact) on the embedding or graph, and not be close to elements of other classes (*i*.*e*. separation or local homogeneity). Their exact meanings will vary across contexts, and we explore how different metrics, some well-established and others novel, relate to these properties.

#### Embedding assessment

Embedding-based metrics are often internal metrics used to evaluate clustering quality, but can also be used as external metrics using the true class labels. A cell embedding can be evaluated either directly using distance measures within the embedding space, or indirectly through secondary information derived from these distances, such as a nearest-neighbor graph. The latter will be discussed in the following section. We discuss three representative embedding metrics: the Silhouette score [31], the composed density between and within clusters index (CDbw) [32], and the Density-Based Clustering Validation index (DBCV) [33]; see Table 2 for details. They are all aggregated metrics aiming to balance compactness and separation. Notably, CDbw consists of three different components—CDbw cohesion, CDbw compactness and CDbw separation—which can be used as property-based evaluations.

**Table 2.**
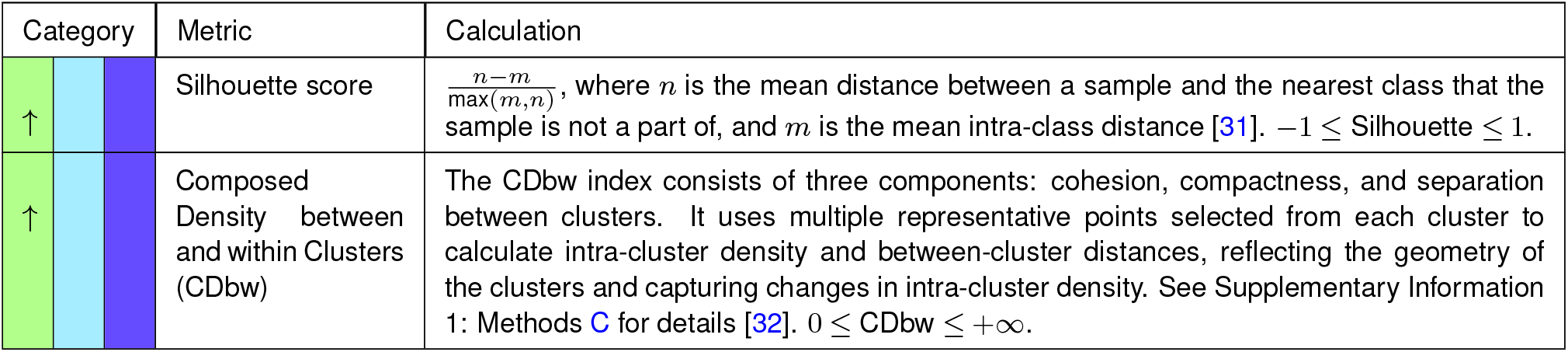
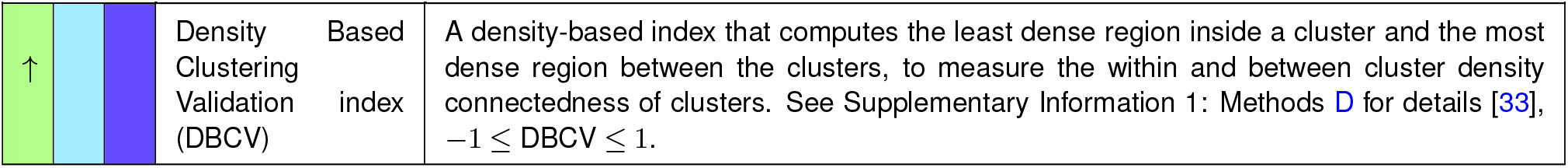
Embedding-based evaluation metrics. Metrics are categorized according to whether they are aggregated or track a specific desirable property, and the minimal unit of calculation (cell, class/cluster, or dataset). In the first column, a ↑ indicates that a higher metric value is better, while a ↓ means that a lower value is prefered. 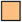 partition-based metric 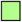 embedding-based metric 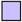 graph-based metric 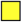 partition-based metric with spatial information 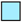 aggregated metric 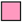 property-based metric 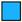 element-level evaluation 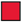 class/cluster-level evaluation 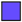 dataset-level evaluation

| Category | Metric | Calculation |
| --- | --- | --- |
| ↑ | Silhouette score | $\frac{n-m}{\max(m,n)}$, where $n$ is the mean distance between a sample and the nearest class that the sample is not a part of, and $m$ is the mean intra-class distance [31]. $-1 \leq \text{Silhouette} \leq 1$. |
| ↑ | Composed Density between and within Clusters (CDBw) | The CDBw index consists of three components: cohesion, compactness, and separation between clusters. It uses multiple representative points selected from each cluster to calculate intra-cluster density and between-cluster distances, reflecting the geometry of the clusters and capturing changes in intra-cluster density. See Supplementary Information 1: Methods C for details [32]. $0 \leq \text{CDBw} \leq +\infty$. |
| ↑ | Density Based Clustering Validation index (DBCV) | A density-based index that computes the least dense region inside a cluster and the most dense region between the clusters, to measure the within and between cluster density connectedness of clusters. See Supplementary Information 1: Methods D for details [33], $-1 \leq \text{DBCV} \leq 1$. |

The Silhouette score is frequently used to evaluate single cell embeddings [3]. At the dataset level, it is a ratio between the separation between classes, understood as the distance between their centroids, and the compactness of the classes, understood as variance around the centroids. As such, it implicitly assumes hyperspherical classes [33, 34]. In contrast, both CDbw and DBCV indices are designed to handle arbitrarily-shaped clusters or classes. CDbw achieves this by representing each class with multiple points instead of a single one. It defines between-class separation through the minimum pairwise distance between representatives, normalized by inter-class density. Compactness, in this case, is defined based on intra-class density, which assesses how tightly the classes’ points are packed around representative points. Additionally, it introduces cohesion to measure the uniformity of intra-class density changes across representatives. DBCV also employs a notion of density, with compactness referring to the density within a class, specifically the maximum distance within the minimum spanning tree of the class, while separation corresponds to the maximum density between two classes.

In Figure 3, we use toy examples to explore how these three metrics behave on classes (clusters) of different shapes. In Figure 3a-b, both scenarios feature two classes sampled from 2D Gaussian distributions. By varying the covariance, the classes take on different shapes, with class A in Scenario 1 forming an elongated cluster, while in Scenario B, it is more globular. In Scenario 1, despite a clear separation between the two classes, many negative Silhouette scores appear due to the elongated shape of class A, resulting in an overall score similar to that of Scenario 2 (Figure 3b), where the classes are visually not as well-separated. For CDbw, although the overall scores are similar between the two scenarios, its components clearly show that the class separation is better in Scenario 1, while compactness and cohesion are worse. This provides a more comprehensive comparison. From a density perspective, DBCV favors Scenario 1 (Figure 3b), likely because the density separation in Scenario 1 is better.

In Figure 3c-d, we highlight two scenarios with moon-shaped classes (*i*.*e*. representing non-convex clusters). Here, density-based clustering methods such as HDBSCAN can correctly separate the two classes, while kmeans clustering, which assumes globular clusters, does not perform well. This performance difference is clearly identified by DBCV and CDbw (Figure 3d), whereas the Silhouette score fails to capture it. Notably, CDbw highlights the separation differences between the two predictions in Figure 3c, but not their compactness differences (independently from the number of representative points used, see Figure S2).

We note that the assumptions about the underlying data structure captured by cell embeddings can vary depending on the downstream applications [35, 36]. For instance, when 2D embeddings are used for visual cluster validation or exploring class relationships—where the global relationships and distances in the embedding space are key [35]—the Silhouette, CDbw, and DBCV may provide valuable insights, as they all operate on the level of cluster relationships and incorporate the notion of distance. However, special caution is needed with the Silhouette score due to its sensitivity to class shapes, since shape can vary significantly between embeddings, and shape distortions introduced by non-linear dimensional reduction methods in single-cell data are common [36].

For downstream applications such as clustering, the assumptions used depend on the analysis workflow. Current best practice for clustering single-cell omics data [1, 37, 38] often involves constructing a kNN/SNN graph and applying graph-based clustering (see Glossary). This assumes that the embedding space preserves distances in local neighborhoods, but not necessarily global distances. Embedding-based metrics still rely on global distances, and because of this difference in assumptions, lower values from embedding-based metrics do not necessarily imply a poorer clustering outcome. Thus, we argue that metrics that focus on local neighborhood relationships, which we introduce in the next section, are better aligned with the clustering workflow. There might however be other downstream tasks for which obtaining more globular class representations would for instance be important, and in such a context the Silhouette score would still be valuable.

Given the demonstrated importance of local similarities in clustering, the most intuitive evaluation at the embedding level would be to examine how well the local neighborhood distances are preserved in the embedding space. However, there is typically no ground truth available for this. For instance, Euclidean distances in the high-dimensional “ambient” space (see Glossary) cannot be assumed to accurately represent the true relationships due to the curse of dimensionality [36, 39]. [40].

Even in a low-dimensional embedding space, there are contexts in which Euclidean distances are inappropriate [40, 41]. Figure 4 shows a single-cell ATAC-seq dataset where a latent space was derived using Latent Semantic Indexing (LSI) (see Supplementary Information 1: Methods B). In Figure 4a, Uniform Manifold Approximation and Projection (UMAP) was performed for visualization using the LSI embedding as input. Figure 4b shows Silhouette scores of true classes using either Euclidean or Cosine distances on the LSI space, colored by the class of the closest centroids. Our observations revealed that for certain cell types (*e*.*g*., vascular smooth muscle cells), despite UMAP’s capability to effectively identify and separate these cells by relying on local connectivity, the Silhouette score using Euclidean distance in the LSI embedding space often resulted in a large proportion of negative scores, while using Cosine distance greatly alleviates this problem. In the scATAC-seq example, this is chiefly because of technical library size variation across cells: the deeper they are sequenced, the more they distinguish themselves from other classes, and hence the more spread out in the embedding (Figure 4c-e), creating Euclidean (but not Cosine) distances between cells that represent the same pattern, but differ in information content.

#### Graph structure assessment

Graph-based clustering has become increasingly popular across multiple fields due to its effectiveness in analyzing complex networks [30, 42], including in single-cell omics data [1, 38, 43]. The general idea of graph-based metrics is to assess whether connections within communities are significantly denser than those with the rest of the network [30]. This assumes that the graph is locally homophilic, *i*.*e*. nodes with similar attributes are more likely to be connected [42, 44], which is guaranteed by the construction of the kNN/SNN graph. In addition, the construction of such graphs only assumes that distances are preserved locally in the embedding space, mitigating problems related to non-Euclidean spaces.

The intuition underlying graph clustering can be linked to the desirable properties previously discussed, where *homogeneity* indicates that connections primarily occur between nodes of the same class, and *compactness* implies that any pair of nodes within a class exhibits expected similarity, reflecting the cohesiveness of the community (Table 3). Metrics for homogeneity are typically computed for the neighborhood of each node, but can be aggregated for evaluation at varying levels of granularity—per class, or across the entire graph. The simplest, most intuitive metric, which we call neighborhood purity (NP), computes the proportion of the neighborhood sharing the node’s class. A diversity metric commonly used in the single-cell field, the Local Inverse Simpson’s Index (LISI) [46, 47], can also be inverted (i.e. the Simpson Index) to evaluate homogeneity. The Proportion of Weakly Connected (PWC) nodes [4] and modularity [45] have also been used (See Table 3). Purity and LISI/SI operate at the node level^1^, PWC at the class level, and modularity at the graph level. Figure 5a shows the behaviour of these metrics using toy examples. While NP effectively measures the difference between graph 1 and the other two, it fails to capture the nuances in homogeneity between graph 2 and 3. Instead, the small increase in LISI from graph 2 to 3 nicely tracks the increased class diversity (and hence decreased homogeneity) among neighborhoods of comparable purity. At the class level, PWC (known as Flake-ODF in Savic and Ivanovic [30]) serves as an indicator of class separability that is especially sensitive to nodes that are at risk of being misclassified, making it a valuable measure of homogeneity (Figure S3a). In a real dataset (from Luo et al. [4]), the “B memory” and “B intermediate” subpopulations are difficult to separate during clustering (UMAP in Figure S3b and heatmap in Figure S3c), which is reflected in their large PWC score (Figure S3d). Notably, PWC remains robust across different neighborhood sizes used for constructing the SNN graph (Figure S3d). Finally, modularity is a common way to track the relationship between intra- and inter-class edges at the graph level. Of note, modularity is not limited to tracking homogeneity: because the relative weights of the edges depend on the degree of their nodes, changes in compactness can influence the modularity score. In practice, however, its focus on existing edges (as opposed to “missing” edges between elements of the same class) means that the score is mostly driven by homogeneity (see Figure 5b, “modularity”).

**Table 3.**
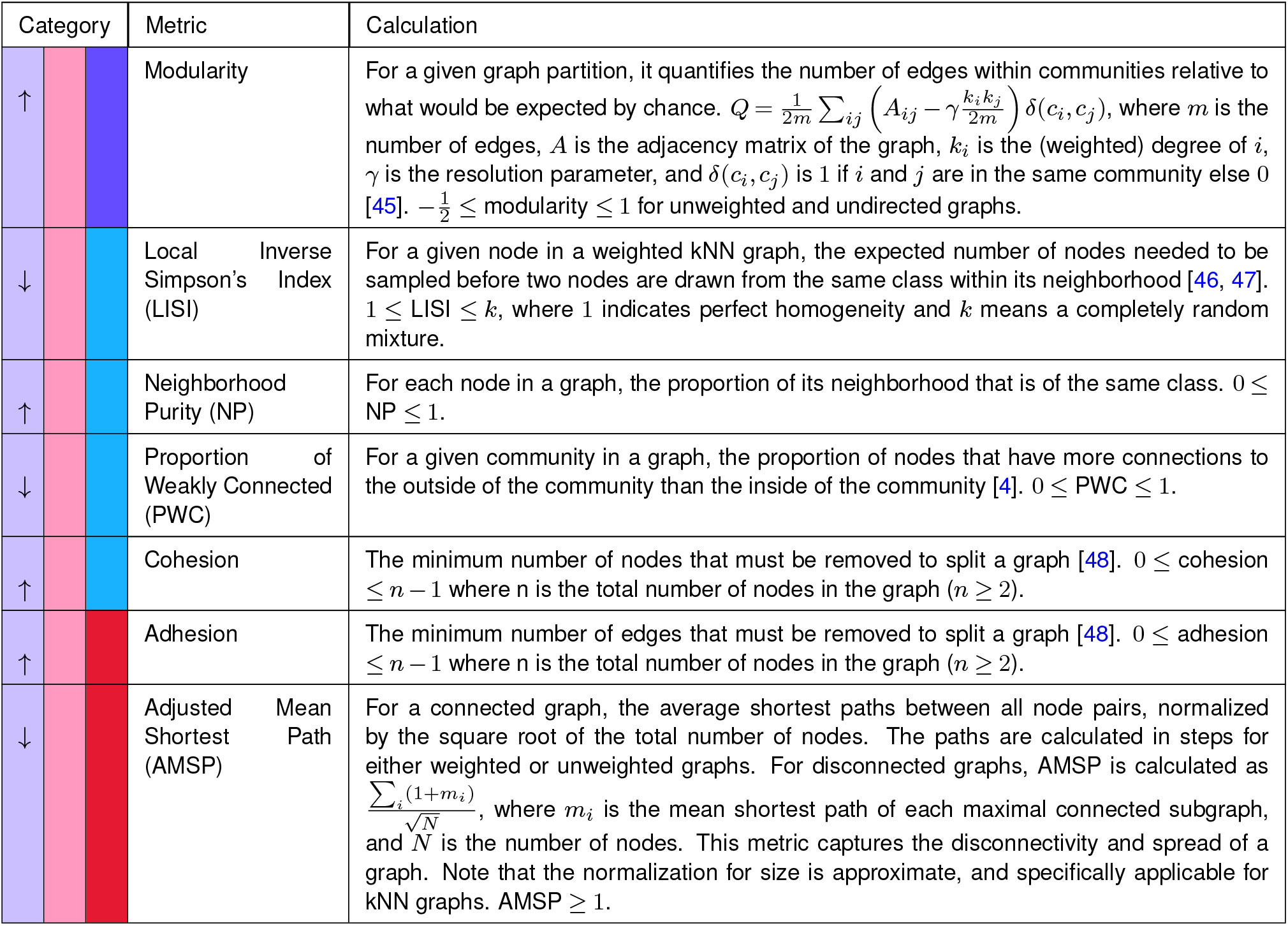
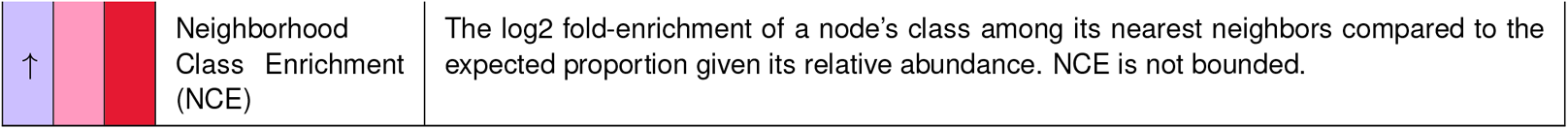
Graph-based evaluation metrics. Metrics are categorized according to whether they are aggregated or track a specific desirable property, and the minimal unit of calculation (cell, class/cluster, or dataset). In the first column, a ↑ indicates that a higher metric value is better, while a ↓ means that a lower value is prefered. 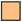 partition-based metric 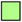 embedding-based metric 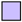 graph-based metric 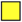 partition-based metric with spatial information 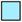 aggregated metric 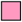 property-based metric 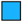 element-level evaluation 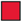 class/cluster-level evaluation 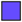 dataset-level evaluation

**Figure 5.**
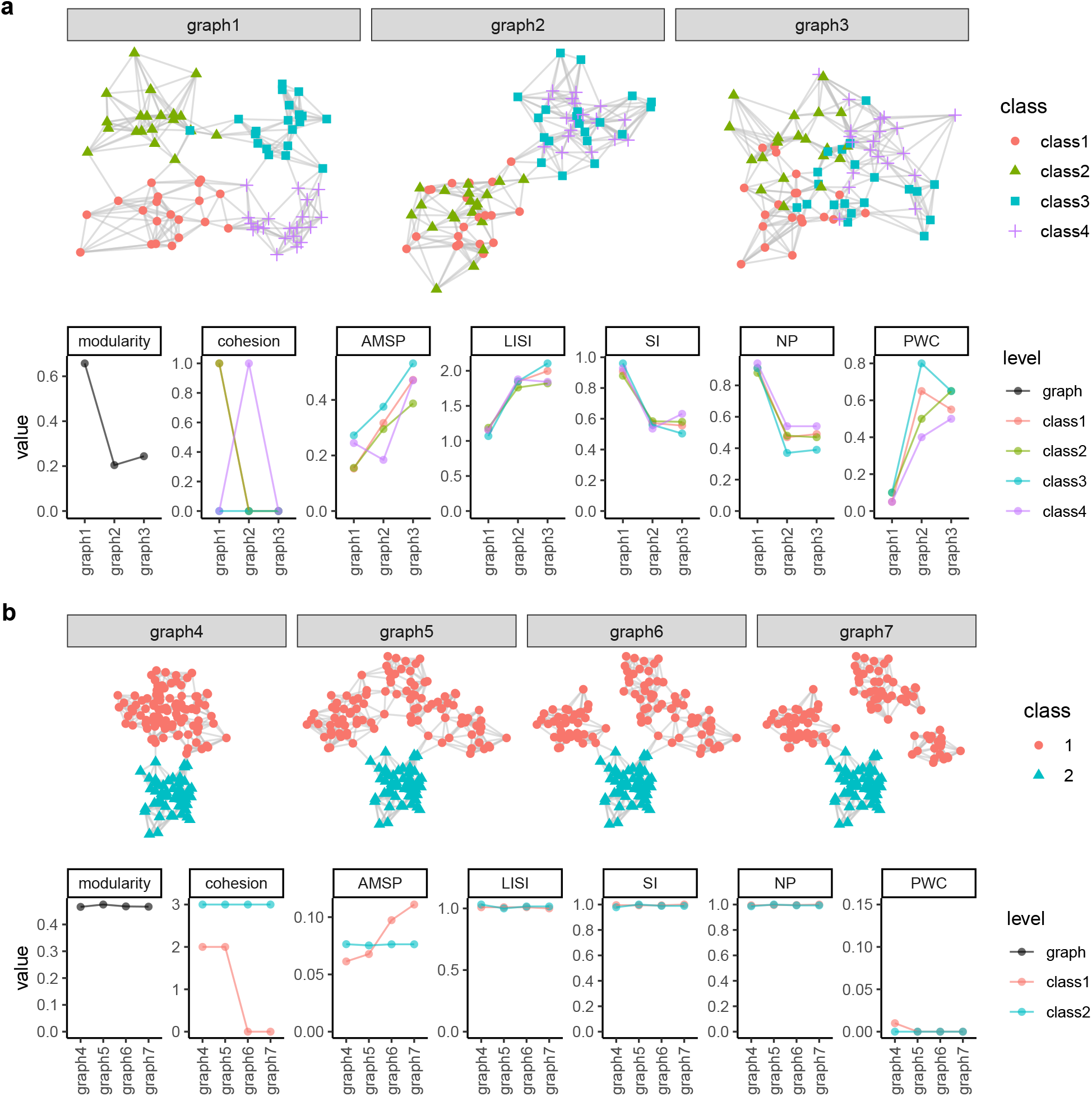
Example graphs and metrics computed on them. **a:** Graphs with varying levels of homogeneity and their corresponding values for different metrics. **b:** Graphs differing in class compactness of class 1, and their corresponding values for different metrics.

Compactness is typically assessed at the class level, with potential to aggregate to the graph level (Table 3). Figure 5b illustrates various graphs that differ in the degree of compactness. While all graphs have a high neighborhood homogenity (as measured by LISI/SI, PWC and NP), graph 4 captures the similarity between nodes of class 1 much better than graph 7 because the subgraph of class 1 is more “compact”; class 2 is identical across all examples. Existing metrics to quantify graph compactness include graph cohesion or adhesion [48], however they are expensive to compute and therefore not scalable to large graphs. Additionally, they are limited to quantifying disconnected class subgraphs because they provide no further distinction once zero cohesion/adhesion is reached, thus making them easily influenced by a few nodes. We propose that a more robust and scalable approach is to look at the distribution of shortest paths between any two nodes for all connected components of the same class, and then aggregate over components. Here, we define a score called “adjusted mean shortest path” (AMSP), which calculates the average shortest paths between all pairs of nodes within each connected components of a class, add 1 to these values to penalize disconnectedness, and then sum across components. Since the mean shortest path in a kNN graph is linearly proportional to the number of vertices (Figure S4), we divide the value by the number of nodes to yield a metric that is less dependent on class size. This nicely captures the differences between graphs in Figure 5b.

Local homogeneity is critical to the quality of clusterings based on nearest neighbors, making the corresponding metrics highly valuable for the evaluation of (single-cell) embeddings and derived graphs. Graph compactness is also important because discontinuities in the graph may cause improper class separation. However, beyond a certain level of compactness (i.e. assuming that the graph is sufficiently connected), there might not be impact anymore on downstream clustering. In practice, the relevance of compactness is further complicated by the issue of ground truth resolution: since real subpopulations may exist within what is labeled as a single class, it might not be desirable to penalize a reduced compactness. Nevertheless, class compactness may be highly relevant in tasks such as meta-cell identification [49, 50], which aim at minimizing within-group variability.

Overall, graph-based metrics are particularly valuable because they avoid some of the biases that affect embedding-based metrics, including global assumptions about distances and class shape. Graph metrics instead focus on the local structure that is critical for common downstream tasks in the single-cell field. They are highly correlated to clustering homogeneity, while avoiding the problem of partition-based metrics that are highly dependent on the clustering resolution. Moreover, graph-based metrics have the advantage of being applicable, and more or less comparable, across various ways of constructing the underlying embedding and calculating distances between cells. Taken together, we believe they are the most useful to evaluate single-cell processing.

#### Comparison of metrics across data representations

To compare the metrics in various realistic contexts, we simulated 2240 datasets in which the number, abundance and differences between classes, as well as their spread is varied; we applied various clustering pipelines, and computed partition-, embedding- and graph-based metrics (see Supplementary Information 1: Methods F). The correlation, across simulations, between pairs of metrics, and their association with specific features of the simulations, are shown in Figure S5). As expected, the major patterns of association between metrics are driven by the property they track, followed by adjustment for chance. Of note, even metrics that are adjusted for chance are significantly associated with the number of clusters called.

### Domain detection in spatial (transcript)omics data

Spatially-resolved molecular measurements enhance single-cell by retaining spatial context, allowing for a more refined examination of tissue heterogeneity and cell subpopulations within physical space. One main task in the various types of spatial omics data (SOD) is the isolation of spatial homogeneous regions defined by molecular, compositional and functional similarities [51, 52]. So-called spatial domain detection, an analog of clustering, results in partitions of cells or spots (we refer to “spots” and array-based examples for simplicity, but the discussion also apply to other types of SOD). These are often compared to manual annotations based on immunohistochemistry and pre-defined markers, which are often taken as ‘ground truth’ [53–56]. In this context, the partition-based metrics discussed earlier can be directly applied to clustering outputs in a spatially-agnostic way. We extend this by applying the same heuristics of desirable properties, and propose new metrics that incorporate spatial information.

#### Non-spatially-aware evaluation

Metrics such as ARI, MI, EH and EC (Table 1) have already been utilized for SOD data for domain detection [53]. Here we focus our discussion on property-based domain-level metrics, including WC, AWC, WH and AWH, highlighting their transparency and interpretability (“domain” could be either a manually-annotated class or an algorithm-derived cluster). We also introduce an element-level metric, Spot-wise Pair Concordance (SPC), which adapts the pair-counting concept to the spot-level, enhancing interpretability, particularly for visualization purposes.

We apply these metrics to the LIBD human dorsolateral pre-frontal cortex spatial transcriptomics data generated with the 10x Visium platform [10]. Figure 6a shows domain detection results across five methods, along with manual annotations; all methods are set to return the “true *k*”, and clusters are matched to classes using the Hungarian algorithm [58]. Figure 6b presents several partition-based metrics comparing the clusters to the manual annotation. The ranking between methods are generally consistent across metrics, except for the differences between precast [59] and CellCharter [15]. The per-domain AWC and AWH in Figure 6c provide deeper insights that are not captured by dataset-level metrics. For example, precast performs worse than CellCharter in preserving white matter (WM), while CellCharter struggles with preserving the layer-like structure in the L3, L4 and L5 classes.

**Figure 6.**
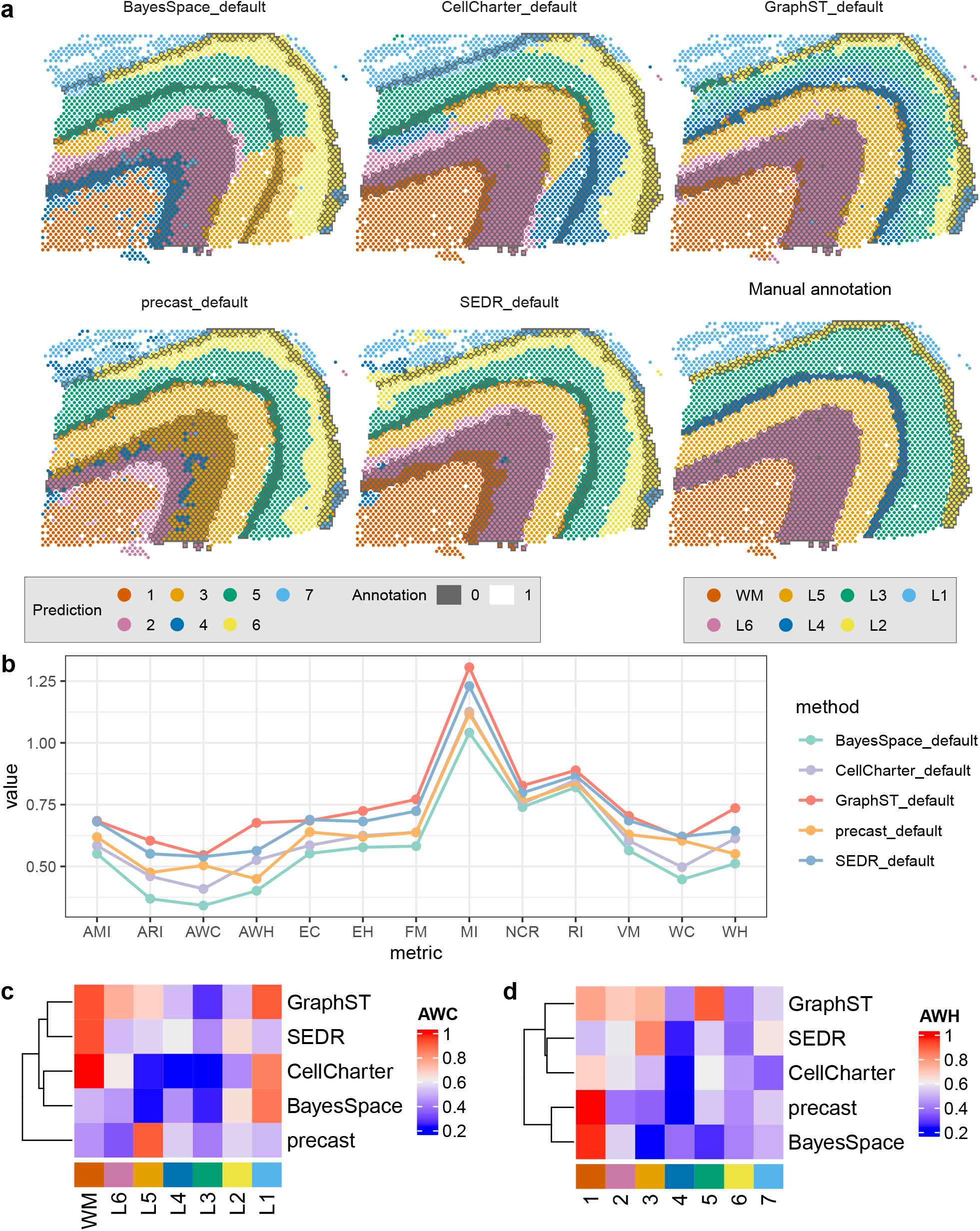
Partition-level metrics can be used for spatial domain detection. The LIBD 10X Visium dataset (slice 151673, Maynard et al. [10]) is used for illustration. In **a**, the last panel shows the manual annotations, while in the remaining panels, each spot is colored according to the prediction from a domain detection algorithm, with colors matched to the corresponding classes in manual annotation. The background is shaded with alternating grey and white regions, representing the annotated adjacent domains from the manual annotation. In **b**, the values of partition-based metrics for each method are calculated. **c**-**d** show the values of AWC (for each class) and AWH (for each cluster).

Furthermore, the domain-level metrics across methods collectively reveal patterns about the manual annotations themselves. For instance, white matter, corresponding to cluster 1, consistently has high AWC and AWH scores, while class L3 shows low completeness scores. This aligns with Figure 6a, where class L3 is frequently split into multiple clusters following the shapes of different cortex layers, suggesting that this domain might exhibit heterogeneity along the axis perpendicular to the cortex layers, possibly representing a gradient layer. Lastly, cluster 4 in Figure 6d appears to be the least homogeneous, and none of the methods clearly identify class L4 of the annotation. Both precast and SEDR [60] treat cluster 4 as a miscellaneous group for spots that may be noisy or contain heterogeneous cell types, while BayesSpace [61], CellCharter, and GraphST [62] label cluster 4 as continuous areas, which are however a mixture of multiple classes based on the ground truth.

#### Spatially-aware evaluation

The metrics above do not use spatial information, which can be used to impose additional constraints. First, adjacent cells are more likely to share common ancestors during morphogenesis and are exposed to similar signaling environments, and therefore to be more similar (*positive spatial autocorrelation*). Second, tissue formation tends to yield continuous structures with relatively smooth boundaries (*spatial continuity*), although the extent of this continuity will vary across tissues. Third, a spot’s identity is shaped not only by its own internal state, but also by its surrounding environment (*spatial contextuality*). Finally, manual annotations of spatial domains, often used as ground truth, can be uncertain [63], particularly near domain interfaces, where the transition between regions may be ambiguous due to spots exhibiting characteristics of multiple domains (*interface uncertainty*); this is especially the case for lower-resolution technologies.

Building on the four characteristics of SOD (above), we propose two pairs of desirable properties. The first pair is *local homogeneity* and *domain continuity*, which respectively estimate the degree of *positive spatial autocorrelation* and *spatial continuity* of SOD. They are internal evaluations, meaning they can be assessed without a ground truth. The second pair includes *neighborhood concordance*, which estimates the extent of concordance between a spot’s spatial context between the predicted and reference partitions, and *interface tolerance*, which tackles *interface uncertainty*. They are external evaluations, which relate to comparisons with a ground truth.

While somewhat intuitive, the notion of domain continuity can have different meanings. On the one hand, it can simply be a higher-level implication of local homogeneity: highly homogeneous local neighborhoods will lead to smooth, continuous domains. On the other hand, there can be continuity in a strict sense with low local heterogeneity (e.g., thin striped pattern). The Spatial Chaos Score (CHAOS) [51], computed at the domain level (Table 4), aims to track the latter, more specific kind of domain continuity. In contrast, two metrics are more focused on local homogeneity: the Entropy-based Local indicator of Spatial Association (ELSA) [57] and the Percentage of Abnormal Spots (PAS) [51] (Table 4). Inspired by Local Indicators of Spatial Association (LISA [64]), ELSA is evaluated at the spot level using categorical variables, but can be aggregated at the domain or dataset level. PAS summarizes at the domain or dataset-level the proportion of spots that are sufficiently inconsistent with their neighborhood (as such, it is analogous to the PWC graph metric). As shown in Figure 7b, such abnormal spots often highlight areas with non-smooth domain boundaries. In contrast, ELSA assigns each spot a continuous value between 0 and 1, where 0 indicates a homogeneous neighborhood (Figure 7d). ELSA highlights boundary regions, with higher values signifying higher diversity. Both PAS and ELSA can be computed across various neighborhood sizes, but are rather robust to it (Figure 7b-d, Figure S6).

**Table 4.**
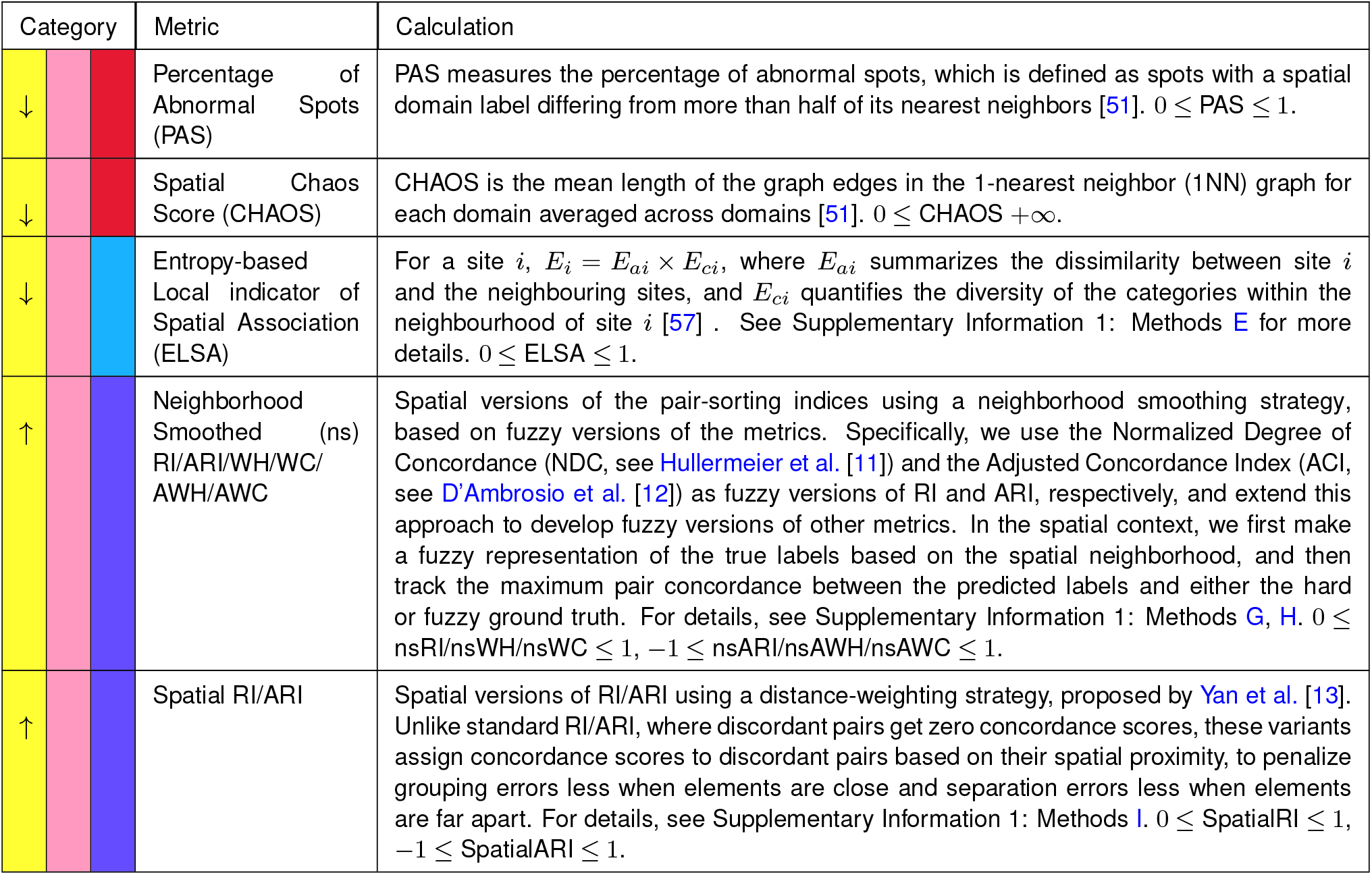
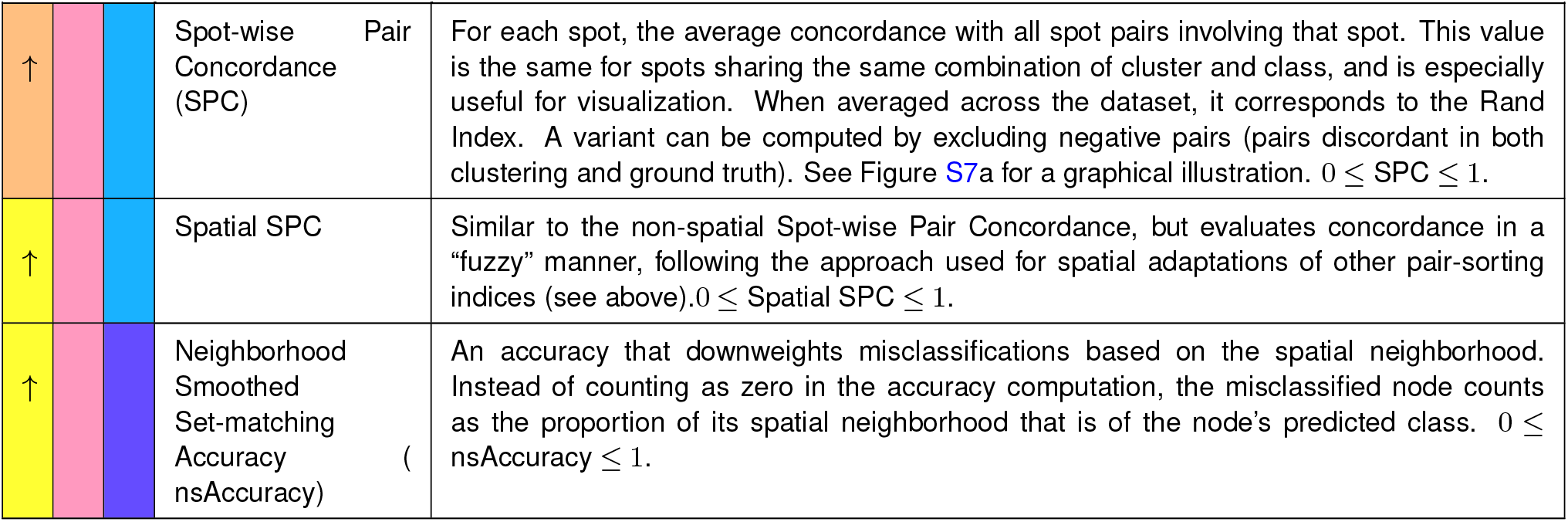
Evaluation metrics for spatial clusterings. Metrics are categorized according to whether they are aggregated or track a specific desirable property, and the minimal unit of calculation (cell, class/cluster, or dataset). In the first column, a ↑ indicates that a higher metric value is better, while a ↓ means that a lower value is prefered 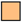 partition-based metric 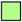embedding-based metric 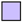graph-based metric 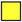partition-based metric with spatial information 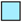aggregated metric 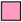property-based metric 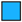element-level evaluation 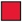class/cluster-level evaluation 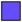dataset-level evaluation

| Category | Metric | Calculation |
| --- | --- | --- |
| ↓ | Percentage of Abnormal Spots (PAS) | PAS measures the percentage of abnormal spots, which is defined as spots with a spatial domain label differing from more than half of its nearest neighbors [51]. $0 \leq \text{PAS} \leq 1$. |
| ↓ | Spatial Chaos Score (CHAOS) | CHAOS is the mean length of the graph edges in the 1-nearest neighbor (1NN) graph for each domain averaged across domains [51]. $0 \leq \text{CHAOS} + \infty$. |
| ↓ | Entropy-based Local indicator of Spatial Association (ELSA) | For a site $i$, $E_i = E_{ai} \times E_{ci}$, where $E_{ai}$ summarizes the dissimilarity between site $i$ and the neighbouring sites, and $E_{ci}$ quantifies the diversity of the categories within the neighbourhood of site $i$ [57]. See Supplementary Information 1: Methods E for more details. $0 \leq \text{ELSA} \leq 1$. |
| ↑ | Neighborhood Smoothed (ns) RI/ARI/WH/WC/AWH/AWC | Spatial versions of the pair-sorting indices using a neighborhood smoothing strategy, based on fuzzy versions of the metrics. Specifically, we use the Normalized Degree of Concordance (NDC, see Hullermeier et al. [11]) and the Adjusted Concordance Index (ACI, see D'Ambrosio et al. [12]) as fuzzy versions of RI and ARI, respectively, and extend this approach to develop fuzzy versions of other metrics. In the spatial context, we first make a fuzzy representation of the true labels based on the spatial neighborhood, and then track the maximum pair concordance between the predicted labels and either the hard or fuzzy ground truth. For details, see Supplementary Information 1: Methods G, H. $0 \leq \text{nsRI/nsWH/nsWC} \leq 1$, $-1 \leq \text{nsARI/nsAWH/nsAWC} \leq 1$. |
| ↑ | Spatial RI/ARI | Spatial versions of RI/ARI using a distance-weighting strategy, proposed by Yan et al. [13]. Unlike standard RI/ARI, where discordant pairs get zero concordance scores, these variants assign concordance scores to discordant pairs based on their spatial proximity, to penalize grouping errors less when elements are close and separation errors less when elements are far apart. For details, see Supplementary Information 1: Methods I. $0 \leq \text{SpatialRI} \leq 1$, $-1 \leq \text{SpatialARI} \leq 1$. |
| ↑ | Spot-wise Pair Concordance (SPC) | For each spot, the average concordance with all spot pairs involving that spot. This value is the same for spots sharing the same combination of cluster and class, and is especially useful for visualization. When averaged across the dataset, it corresponds to the Rand Index. A variant can be computed by excluding negative pairs (pairs discordant in both clustering and ground truth). See Figure S7a for a graphical illustration. $0 \leq \text{SPC} \leq 1$. |
| ↑ | Spatial SPC | Similar to the non-spatial Spot-wise Pair Concordance, but evaluates concordance in a “fuzzy” manner, following the approach used for spatial adaptations of other pair-sorting indices (see above). $0 \leq \text{Spatial SPC} \leq 1$. |
| ↑ | Neighborhood Smoothed Set-matching Accuracy (nsAccuracy) | An accuracy that downweights misclassifications based on the spatial neighborhood. Instead of counting as zero in the accuracy computation, the misclassified node counts as the proportion of its spatial neighborhood that is of the node’s predicted class. $0 \leq \text{nsAccuracy} \leq 1$. |

**Figure 7.**
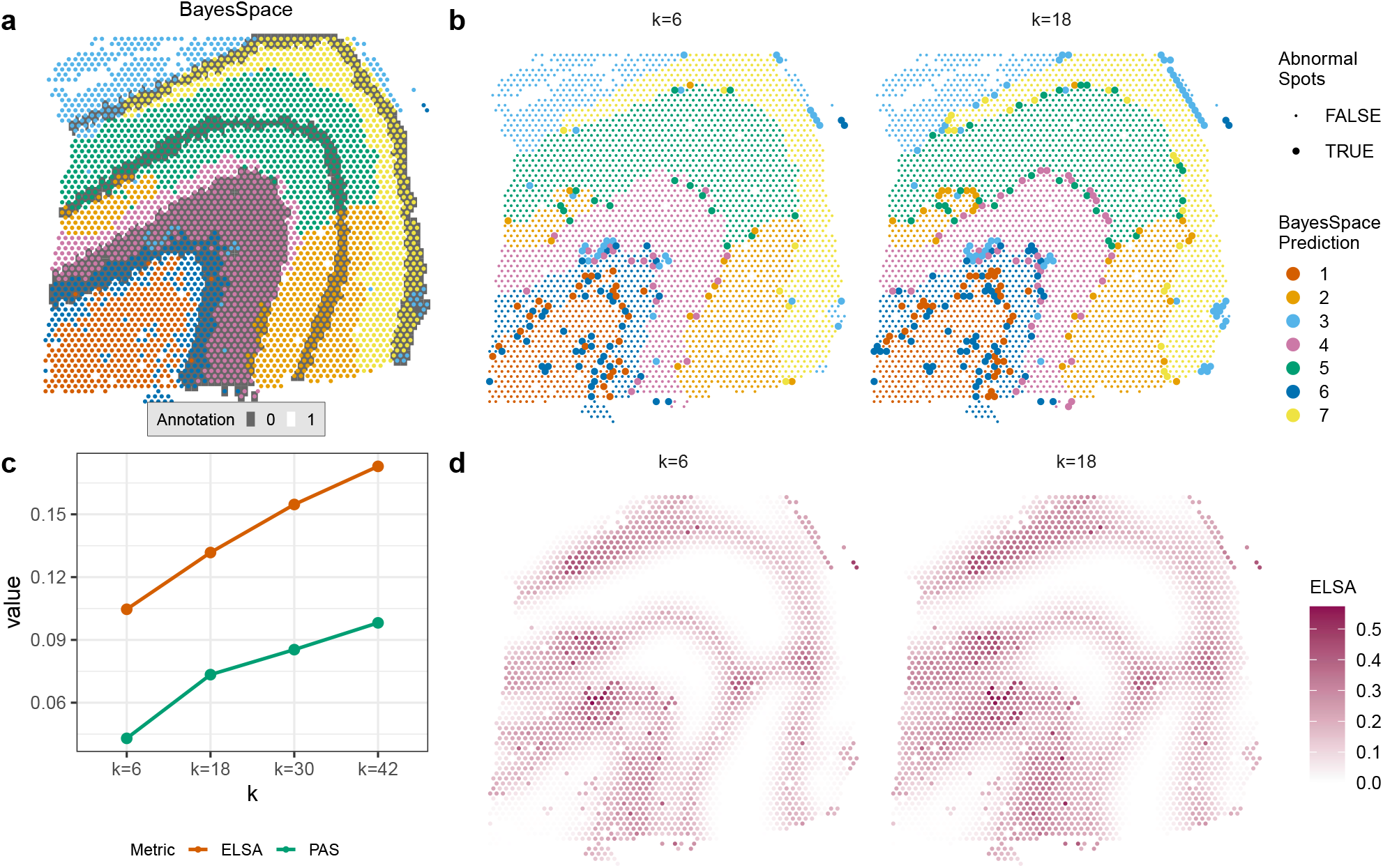
Metrics for spatial clustering evaluation and visualization. The same LIBD 10X Visium dataset as in Figure 6 is used. **a** is identical to the BayesSpace subpanel in Figure 6a, where the border is colored by the predicted domain, and the background is shaded using the ground truth. In **b**, spot-level metric values related to PAS are shown, indicating whether a spot is classified as abnormal or not. **c** displays how the dataset-level ELSA and PAS score change as the size of the neighbourhood (*k*) varies. **d** shows element-level ELSA values. The same visualization for larger *k* are shown in Figure S6.

We also generated toy examples in Figure 8a to investigate further the various metrics. For internal evaluation (Figure 8b), all three metrics correctly identify P3 and P4 as less smooth than P1 or P2, albeit with different dynamic ranges. While CHAOS gives the same scores to P1 and P2 due to the lack of discontinuity, PAS and ELSA consider P1 smoother, as expected. P3 and P4 differ only in the spacing between segments of the blue domain. This leads to a slight increase in CHAOS for P4, reflecting greater spatial discontinuity. From a local homogeneity perspective, ELSA shows a small decrease for P4, indicating a marginal gain in neighborhood consistency. PAS, however, remains unchanged between P3 and P4, as the number of boundary-disrupting spots stays similar. Because these metrics are ground-truth agnostic, the pattern is similar when comparing P5 and P6.

**Figure 8.**
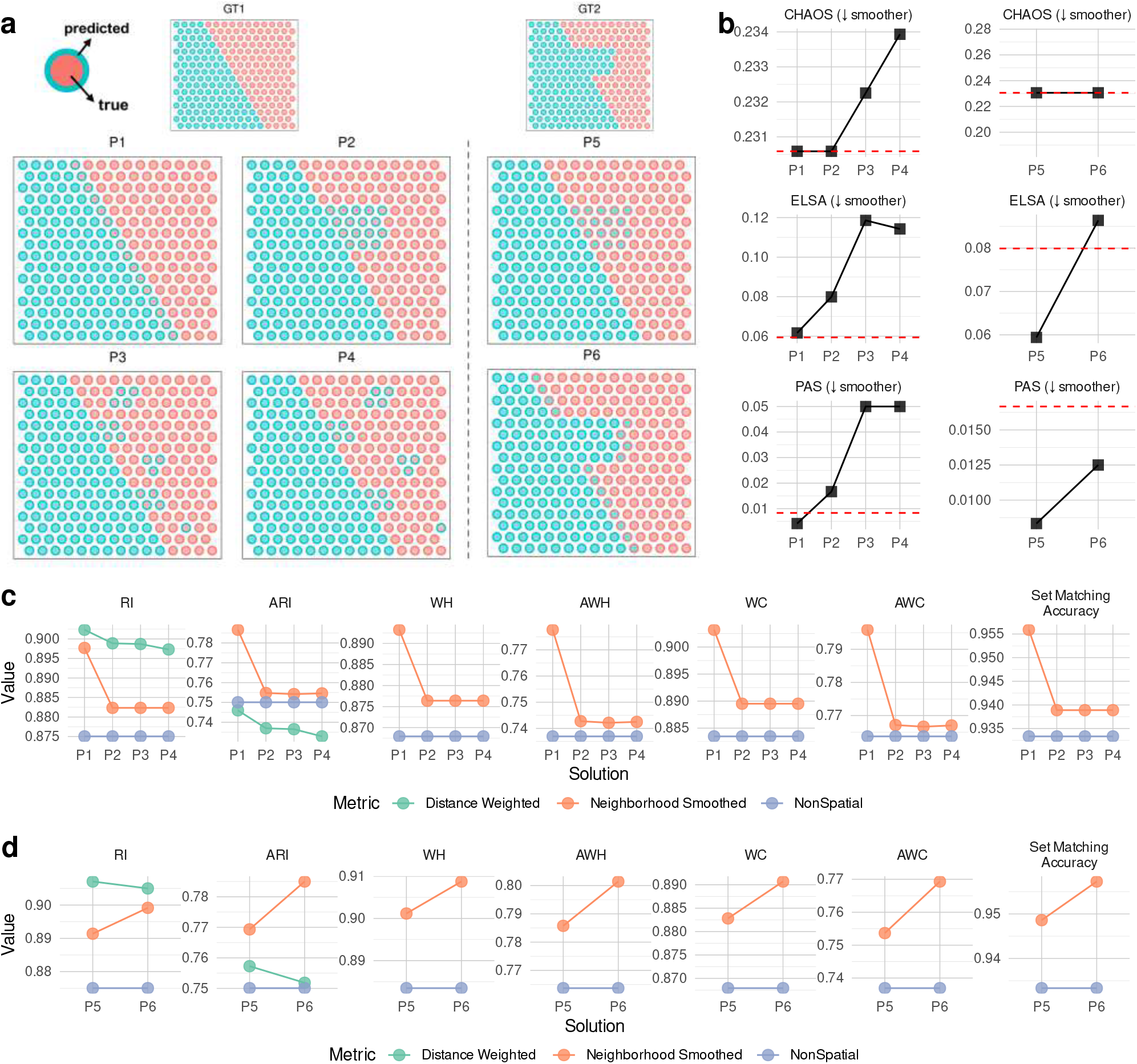
Examples of desirable properties for domain detection, and how different metrics reflect each property. In **a**, the colors within each spot represent the ground truth labels, while the colors around the spot borders indicate the predicted labels. GT1 and GT2 denote two datasets colored by the ground truth domain partition, with P1-P4 showing hypothetical domain detection results based on GT1, and P5 and P6 based on GT2. The number of misclassified spots remains the same across P1 to P3, and also across P5 and P6, but their spatial distribution varies. **b**-**d** demonstrate how each metric compares P1 with P2 and P3, and compares P5 with P6, respectively. **b** shows internal metrics (red dashed lines indicate the values for GT1 and GT2), while **c** and **d** are external metrics. Non-spatial, distance-weighted, and neighborhood smoothed versions are computed for RI and ARI. For the other metrics, non-spatial and neighborhood smoothed variants are included.

At the time of initial writing, there was no existing external partition-based metrics that take the spatial information into account. To include this spatial dimension into partition-based evaluation, we developed two strategies.

The first approach is based on set matching (see Glossary), and evaluates the agreement between a spot’s predicted label and the true labels of both the spot and its neighborhood, capturing spatial relationships. We refer to this as the neighborhood smoothed set-matching accuracy (nsAccuracy, see Table 4). As shown in Figure 8c and d, nsAccuracy correctly reflects the property of interface tolerance, as misclassified spots at the domain boundary are assigned to the adjacent domain, reducing the penalties for such spots. Since misclassifications in P1 are located more at the interface between domains that those of P2, P3 and P4, it has higher set-matching accuracy. P2, P3 and P4 further demonstrate that when the number of interface misclassifications is the same, nsAccuracy remains consistent, regardless of the distribution of other misclassifications within the domain. This confirms that nsAccuracy effectively captures interface tolerance in a desirable way, while not expecting more smoothness than is present in the ground truth (Figure 8c-d). In contrast, internal metrics in Figure 8b assign P5 a smoother score than P6, ignoring the non-smooth nature of GT2. Despite being simple and easy to interpret, this nsAccuracy has two major limitations: it requires matching clusters to classes, which can be ambiguous, and it is not adjusted for chance (i.e. expected agreement based on random allocations).

Another strategy we propose is to make either or both the clustering and the ground truth fuzzy (see Glossary), by averaging a spot’s identity with that of its neighborhood, and then employ metrics for fuzzy clustering evaluation [11, 12]. Building on the permutation-based fuzzy version of the ARI [12], we developed a fuzzy version of the (adjusted) Wallace indices (*i*.*e*. WH/AWH and WC/AWC (see Table 4). Based on this, we defined ‘neighborhood smoothed’ versions of the pair-counting metrics (Figure 8c-d and Supplementary Information 1: Methods H). As shown in Figure 8c, these spatially-aware metrics all correctly prefer P1 to P2-P4, and P6 to P5, thus displaying interface tolerance.

To better understand the metric behavior with fuzzy representations, let us look at domain-level metric values. When interpreting SpatialWC and SpatialAWC in Figure 8, one should consider the values for each *ground-truth* domain (*i*.*e*., class), as shown in Figure S9. In Figure 8a, for spots of the red class, although P1-P4 misclassify the same number of spots into the blue cluster, the completeness of the red class in P1 is higher than in P2-P4. Because there are more misclassified spots in P1 that are from the interface of red and blue classes that contain partial blue memberships, the error is considered less severe. On the contrary, when interpreting SpatialWH and SpatialAWH in Figure 8, one should consider the values for each *predicted* domain (*i*.*e*., cluster), as shown in Figure S9. In Figure 8a, for spots of the blue cluster, although P1-P4 incorrectly mix the same number of spots from the red class into the blue cluster, these red spots in P1 are all on the interface between the blue and red classes in GT1, and thus have partial membership of the blue class, making such errors less penalized than the errors in P2 -P4. For P2 -P4, the number of misclassified spots at the class interface are identical, and thus the spatial metrics have identical values (within the range of variation of the permutations for adjusted metrics).

During the writing of this research, Yan et al. [13] proposed a different set of spatially-sensitive properties. They stipulated that: 1) incorrectly grouping two elements (e.g. spots) into the same domain should be penalized less if the elements are close to each other; and, 2) incorrectly separating two elements should be penalized less if they are distant from each other. Rather than tackling *interface uncertainty*, as do the aformentioned fuzzy metrics, these properties emphasize *positive spatial autocorrelation* and *spatial continuity*, both of which are expected in SOD. The authors developed a spatially-aware version of the Rand Index (Spatial RI) by giving discordant pair a non-zero concordance value based on their spatial proximity. They additionally developed a version adjusted for chance (Spatial ARI). While the proposal and metrics are sound, we found that the original weight functions proposed by the authors do not change sufficiently with distance (see Figure S10), and therefore adjusted them. These ‘distance-weighted’ RI/ARI are shown for our toy example in (Figure 8c-d). Both the distance-weighted and neighborhood smoothed RI/ARI show a decrease in P2-P4 compared to P1, with the strongest decrease for the neighborhood smoothed scores (Figure 8c). They differ, however, in whether P4 is worse than P2 and P3. Neighborhood smoothed RI/ARI stay invariant across P2 to P4, as they are sensitive only to the number of edge errors, not their spatial distribution. In contrast, distance-weighted RI/ARI further decrease in P4, reflecting the greater spatial separation between mislabeled spots and the true blue domain. Which behavior is more desirable depends on assumptions and downstream application. Similarly, the two spatially-aware variants disagree on which of P5 and P6 is worse. The intuition of *edge tolerance* explains why the neighborhood smoothed RI/ARI assign a higher score to P6 than P5, while it is less intuitive why the distance-weighted approach would assign a higher score to P5.

Finally, for element-wise external evaluation and visualization, we developed the Spot-wise Pair Concordance score (SPC) (Table 4). It can incorporate either hard-hard or fuzzy-hard representations of the truth and prediction, making it flexible for either non-spatially-aware or spatially-aware evaluation. This is useful to spatially visualize the different degrees of disagreement between a predicted domain partition and a ground truth without the need for set matching (Figure S7). To facilitate exploration of the aforementioned metrics, we developed a shiny app, available at https://plger.shinyapps.io/poem/, which allows users to visualize results interactively and interpret the behavior of the metrics on customizable toy examples.

### Time and memory complexity

With the emergence of atlas-scale single-cell datasets, it is important to have evaluation metrics that are both time- and memory-efficient. For metrics newly proposed or implemented by our package `poem`, we monitored CPU time and peak memory usage across four dataset sizes by subsampling a large dataset (Figure S11, Figure S12). With the exception of DBCV, all tested partition-, embedding-, and graph-based metrics demonstrate high scalability. DBCV is computationally intensive in both time and memory due to the need to compute pairwise distance matrices and minimum spanning trees within each cluster.

Among spatial metrics using distance-weighting, the original implementation of SpatialARI [13] by Yan et al. consumes excessive memory and fails with an out-of-memory error for the dataset with 136,870 cells. In contrast, our optimized version significantly reduces memory usage (by 10- to 100-fold) at comparable runtime. Spatial metrics using neighborhood smoothing are comparatively slow but remain memory-efficient.

## Discussion

Evaluation metrics play a crucial role in shaping our understanding of the strengths and weaknesses of computational methods, guiding their development, and ultimately influencing our interpretation of biology. However, the current practice of selecting evaluation metrics in the single-cell omics field often lacks a clear and rational basis. In this work, we leverage the concept of desirable properties to gain a deeper understanding of the behavior of each metric, offering a structured guideline for rational metric selection.

To illustrate and explore these desirable properties, we make extensive use of carefully designed toy examples. These examples, while simple, offer clear insights that can be difficult to extract from real-world datasets, where multiple factors are often entangled. Despite their simplicity, these examples demonstrate what does or does not influence a metric, thereby helping the interpretation in a way that extends to real-world datasets.

In single-cell -omics, the identification of rare subpopulations is often most challenging, and some of the most widely used metrics, such as the ARI, are less sensitive to misclassification errors happening in smaller populations. They are also, we argue, less interpretable. Given that each metric captures different desirable properties and functions at different evaluation levels, we generally advocate the use of property-based metrics and those that operate at finer levels, such as class- or spot-level metrics, for fine-grained method comparison. As demonstrated in the clustering examples and domain detection examples, they offer greater transparency and interpretability compared to aggregated, dataset-level metrics.

We extend the most relevant desirable properties from partitions to embeddings to graphs, enabling a similar and structured evaluation across these levels, and emphasize that desirable properties should be carefully considered within the biological context of a study. By examining the relationships and trade-offs between different properties, we aim to help researchers select metrics tailored to their biological questions and the complexities of different datasets. This approach aids in developing useful metrics, as well as in addressing the challenge of interpreting benchmarking results that involve multiple metrics and datasets with inconsistent rankings of methods.

By understanding metrics from the perspective of desirable properties, researchers can more precisely assess the validity of the underlying assumptions. In our discussion, we advocate evaluating single-cell data from a connectivity standpoint at the graph level, as it requires fewer global assumptions than embedding-based evaluations, and has proven to be highly effective in capturing statistical patterns in single-cell datasets. In addition, it is preferable over partition-based evaluation because it mitigates issues caused by misaligned levels of biological granularity between the prediction and ground truth. To facilitate this practice, we introduce novel metrics–PWC and AMSP–that align with this perspective. In particular, we introduce AMSP as a measure of graph compactness: while compactness is not always crucial when evaluating the discrimination of cell populations (because the ground truth annotation typically does not exclude the existence of unknown sub-populations), it can be for other tasks. For example, compactness is critical to accurate trajectory inference, where discontinuities could lead to wrong inferences. An implementation of all of the metrics discussed here (and more), including the novel metrics and fuzzy adaptations, are available in our R package poem (https://bioconductor.org/packages/poem).

In addition, we extend our work to the problem of spatial domain detection in SOD. The evaluation of domain detection methods has often relied on internal metrics or external metrics that do not account for spatial relationships. To address this, we developed spatially-aware metrics for external evaluation. While we tested our external metrics primarily on array-based SOD, in principle they are applicable to non-array-based SRT datasets by defining neighborhood structures using physical distances between cell centers or boundaries.

Overall, our work emphasizes the contribution of a systematic approach to evaluation metrics grounded in desirable properties. This method prioritizes biological intuition and concerns, rather than uncritically adopting metrics from other fields. In our analysis of spatial domain detection, we demonstrate how different data modalities introduce additional biological considerations, from which we derive new desirable properties. This showcases the strength of our framework not only in providing guidelines but also in generating new principles and inspiring the development of novel metrics.

## Glossary

Partition: a grouping of a set of elements into non-empty subsets, such that every element is included in exactly one subset.
Metric: a standard measure for quantification and evaluation, not to be confused with distance metrics in mathematics and geometry.
External clustering metrics: metrics that assess how well the clustering results match a predefined (e.g. ground-truth) partition of the data.
Adjusting for chance: In clustering, some degree of concordance between two partitions is expected purely by chance. Chance adjustment removes expected random concordance, ensuring that a completely random labeling yields a constant baseline value.
Meta-cell identification: A single-cell omics analysis procedure that groups similar cells and uses their averaged profiles for downstream tasks, usually to reduce noise.
“Ambient” space: here it refers to the log-normalized count matrices filtered for HVGs in scRNA-seq data, or the fragment count matrices after genomic region selection in scATAC-seq data.
Embedding space: the low-dimensional space that cells are projected into (*e*.*g*. the PCA space in scRNA-seq data).
Graph-based clustering: A procedure that represents a dataset as a graph, with invididual observations as nodes connected based on their similarities, community detection algorithms (*e*.*g*. Louvain or Leiden) are used to divide the graph into clusters.
k-Nearest Neighbors (kNN) Graph: A graph that connect nodes to its k most similar neighbors based on a distance measure.
Shared Nearest Neighbors (SNN) Graph: A graph connecting nodes by the extent to which they share their k nearest neighbors.
Set matching indices: Metrics that first solve the optimal matching problem to match or pair each cluster to a class, then calculate some evaluation measures based on that.
Fuzzy clustering: Clustering where a data point can belong to more than one clusters with different degrees of membership.

## Acknowledgements

We wish to acknowledge Robinson Lab members for fruitful discussion and the core SpaceHack 2023 team (https://spatialhackathon.github.io/past.html), especially Jieran Sun, Peiying Cai, and Sarusan Kathirchelvan, for discussions and sharing partitions from methods run on the LIBD dataset.

## Author contributions

SL, PLG, and MDR conceptualized the study. SL and PLG curated and simulated the data. SL and PLG designed and conducted the analysis, developed the metrics and the package, and created the visualizations. MDR and FvM secured funding for the project. MDR provided the primary supervision, while FvM offered supporting advision. SL and PLG drafted the original manuscript with input from MDR. SL, PLG, and MDR collaboratively edited and reviewed the manuscript. All authors approved the final version for publication.

## Code availability

All the metrics we mentioned are implemented in our R package poem (https://bioconductor.org/packages/poem, snapshot on Zenodo: https://doi.org/10.5281/zenodo.15853802). Notebooks, R scripts, and supporting data used for reproducing the analysis and generating the visualizations in this manuscript are available at https://github.com/RoseYuan/metric_paper (snapshot on Zenodo: https://doi.org/10.5281/zenodo.15853790).

## Supplementary Figures

**Figure S1.**
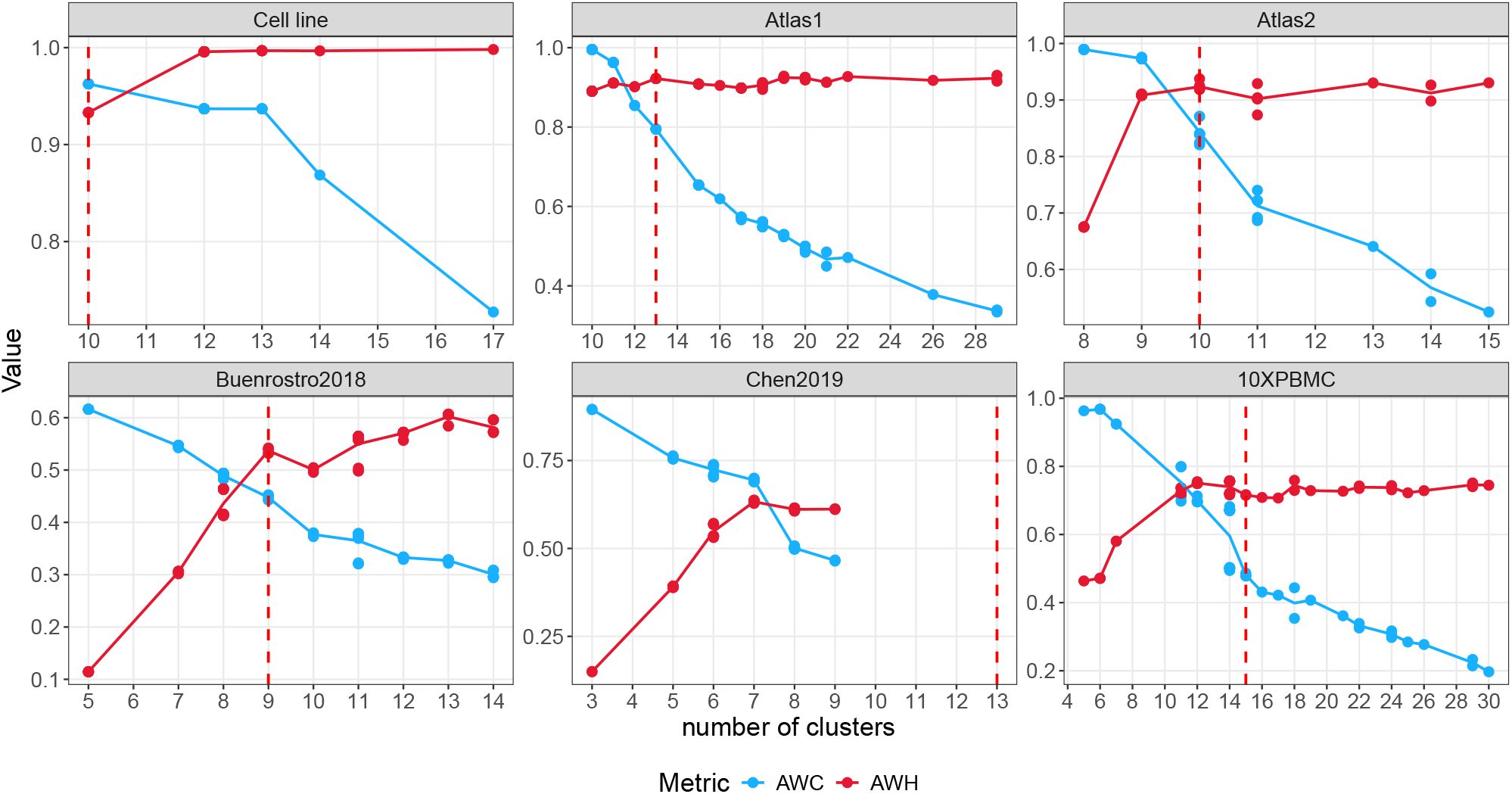
The trade-off between AWH and AWC as the number of clusters varies. Here six single-cell ATAC-seq datasets are used for benchmarking clustering. The red dash line is the number of classes of the ground truth. Each dot is a clustering solution. Different numbers of ATAC clusters are achieved by applying a range of resolutions during Leiden clustering. See [1] for more details.

**Figure S2.**
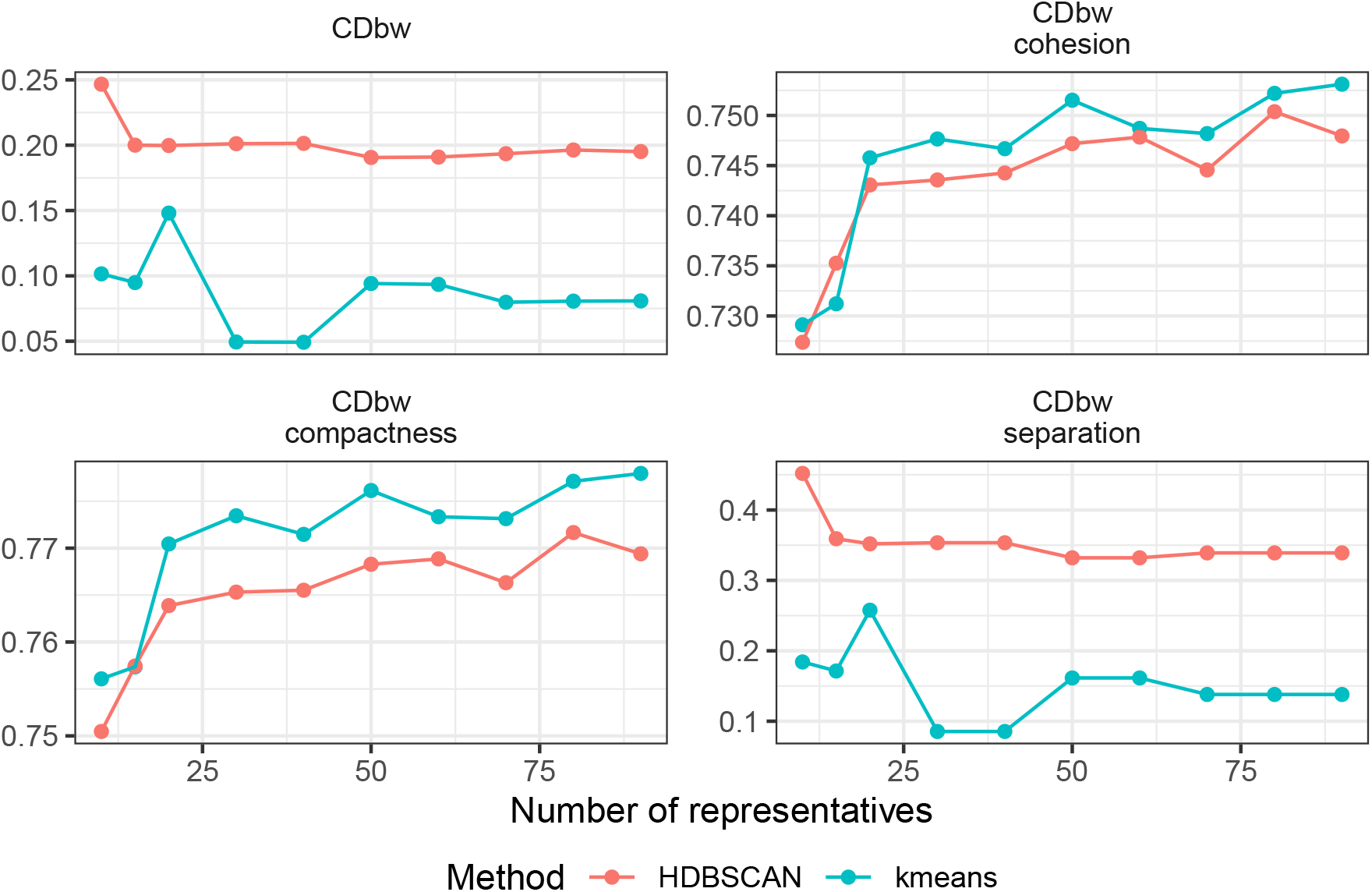
How the number of representative points influence CDbw metrics. In relation to Figure 3 c-d, we calculate the CDbw metrics using a variety of representative numbers. Similar to Figure 3 d where 10 representatives are used, the CDbw and CDbw separation metrics indicate that HDBSCAN produces a better clustering than kmeans for this toy example. However, the CDbw cohesion and compactness metrics fail to capture this distinction and rank them oppositely.

**Figure S3.**
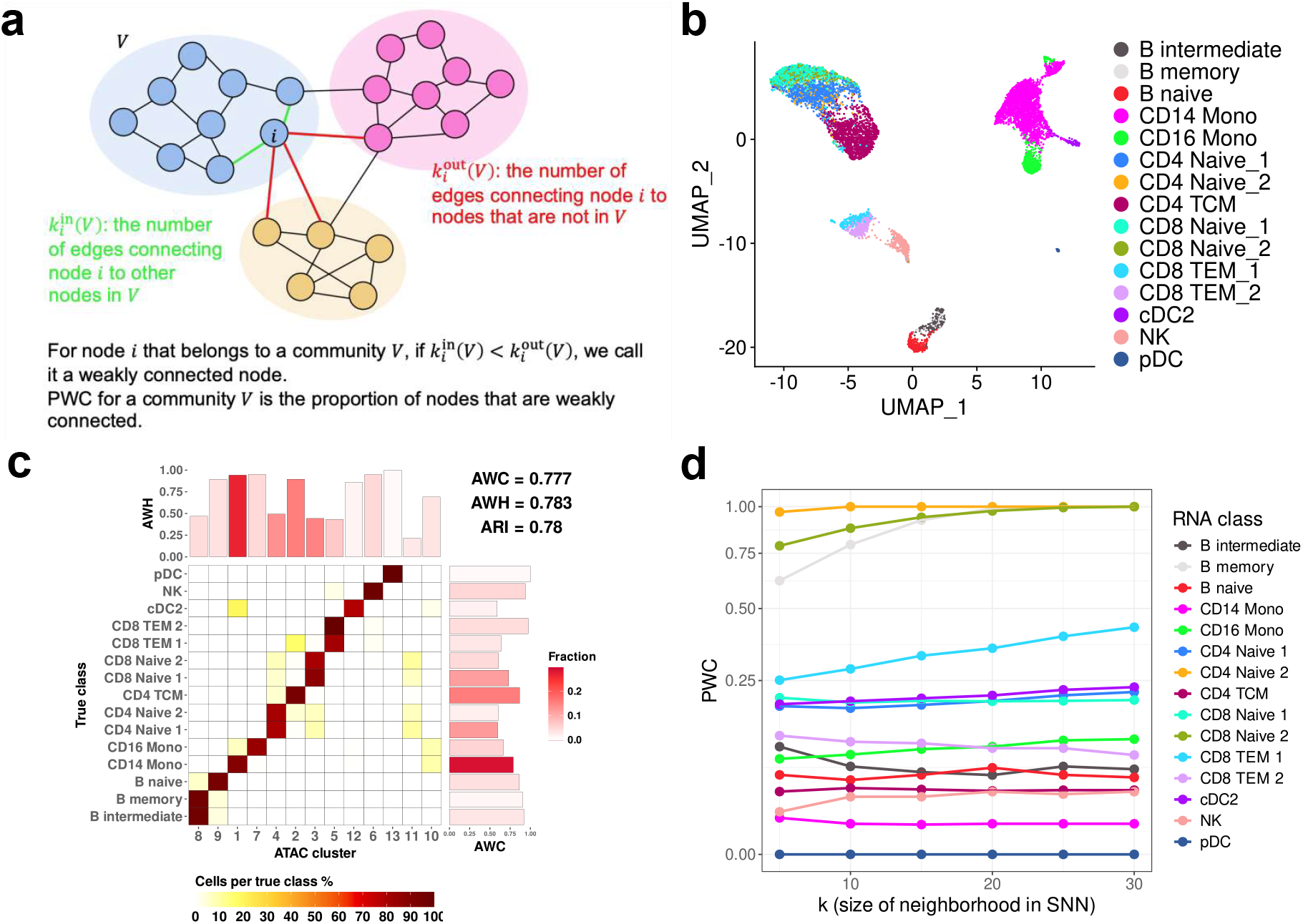
Illustration of the utility of PWC in single-cell datasets. **a** Schemetic overview of how PWC is calculated. **b** The UMAP of this dataset. **c** An example clustering results showing the classes that are failed to be seperated. **d** The value of PWC across a range of k when building the SNN graph.

**Figure S4.**
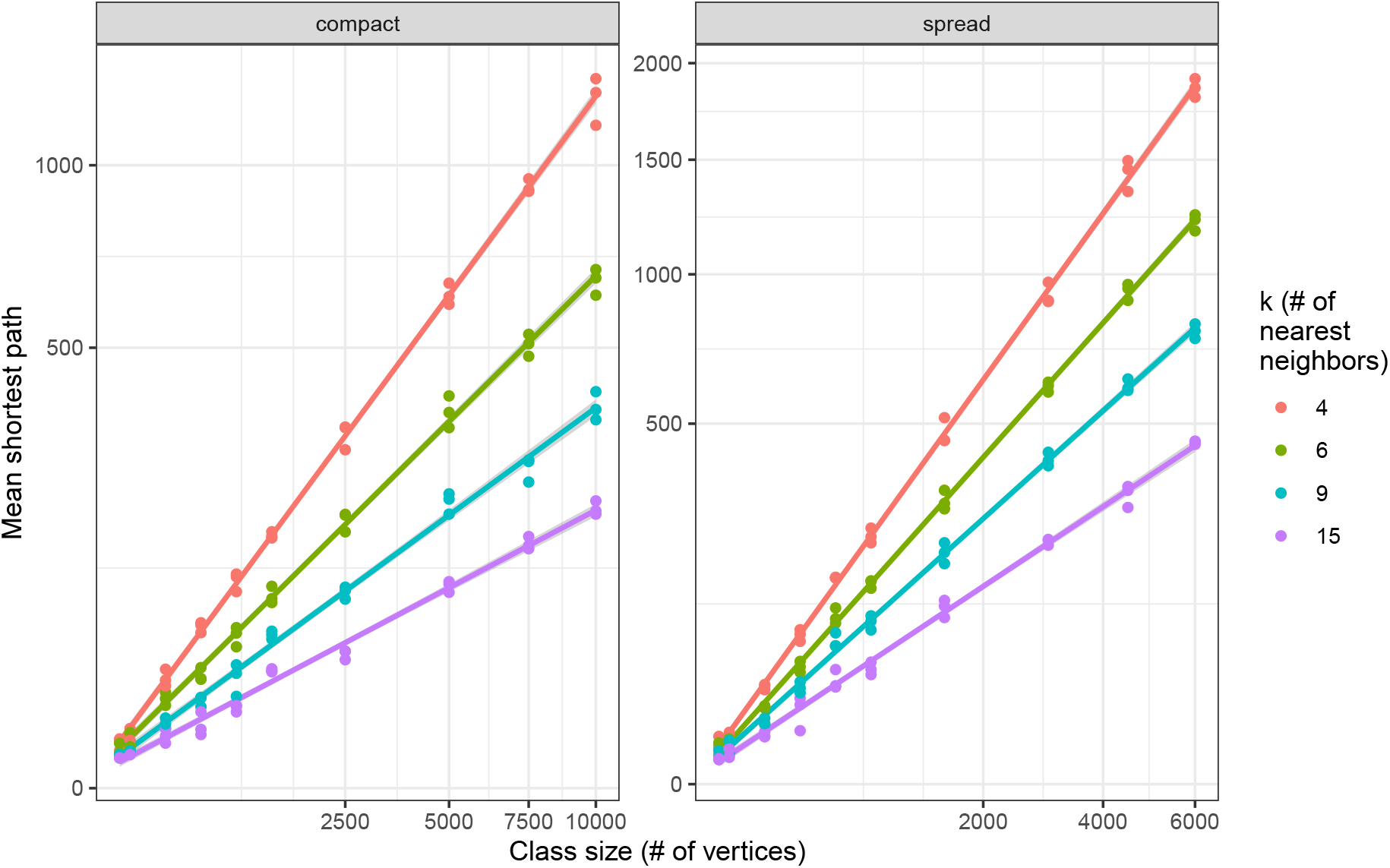
The mean shortest path (in steps) is linearly proportional to the number of vertices. Shown are simulated two-class graphs containing a more compact and more spread class, simulated with the same parameters except for a size factors. Each size is simulated with 3 random seeds.

**Figure S5.**
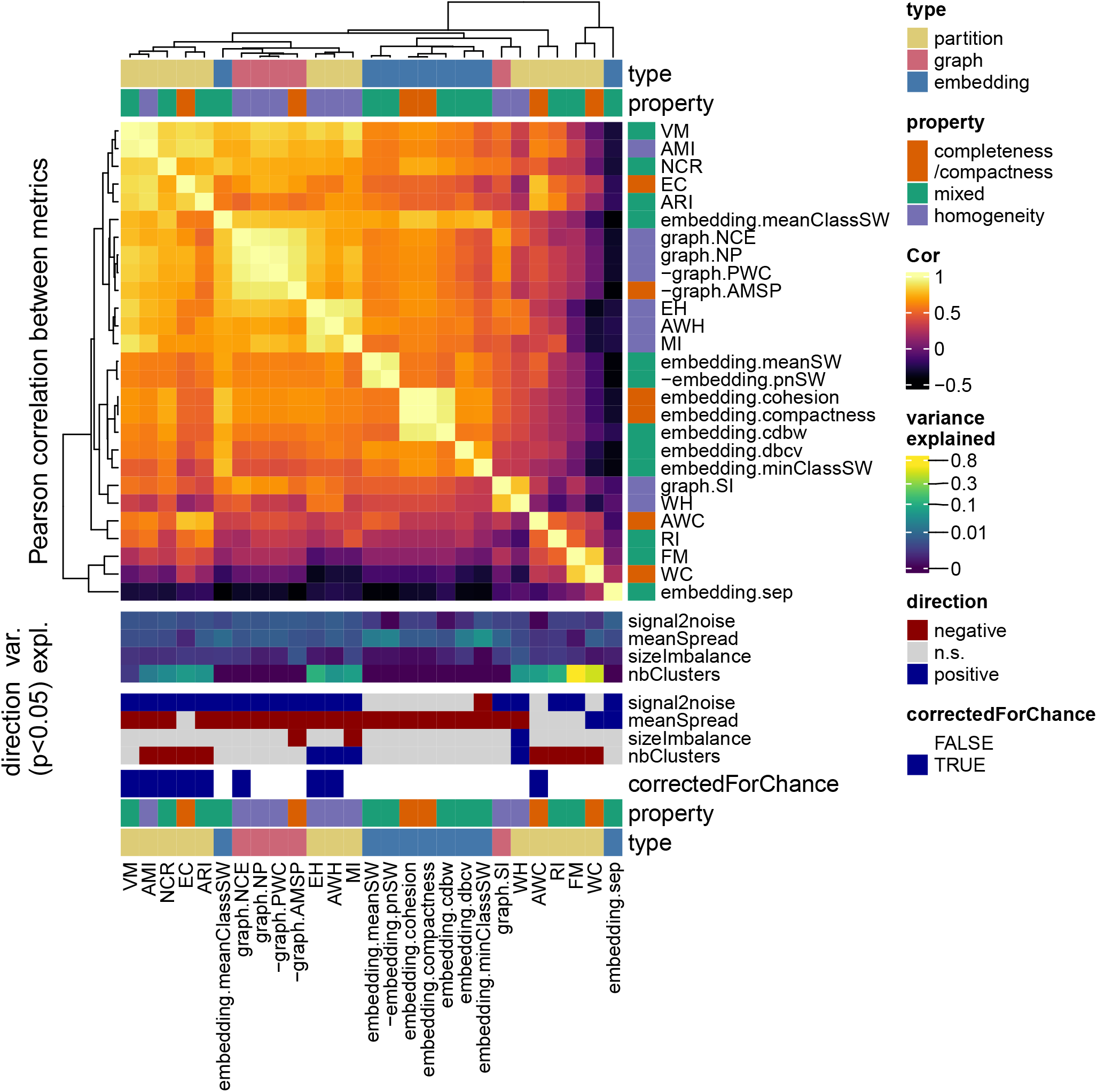
Comparison of metrics across data representations. As explained in Supplementary Information 1, 2240 simulated datasets of 3 or 4 classes with varying abundance, variability, and inter-class distances are used. The main heatmap shows the Pearson’s correlation coefficients between metrics. The bottom heatmaps are the percentage of variance explained by each simulation factor using a linear regression model, the effect directions, and the metric categories, respectively.

**Figure S6.**
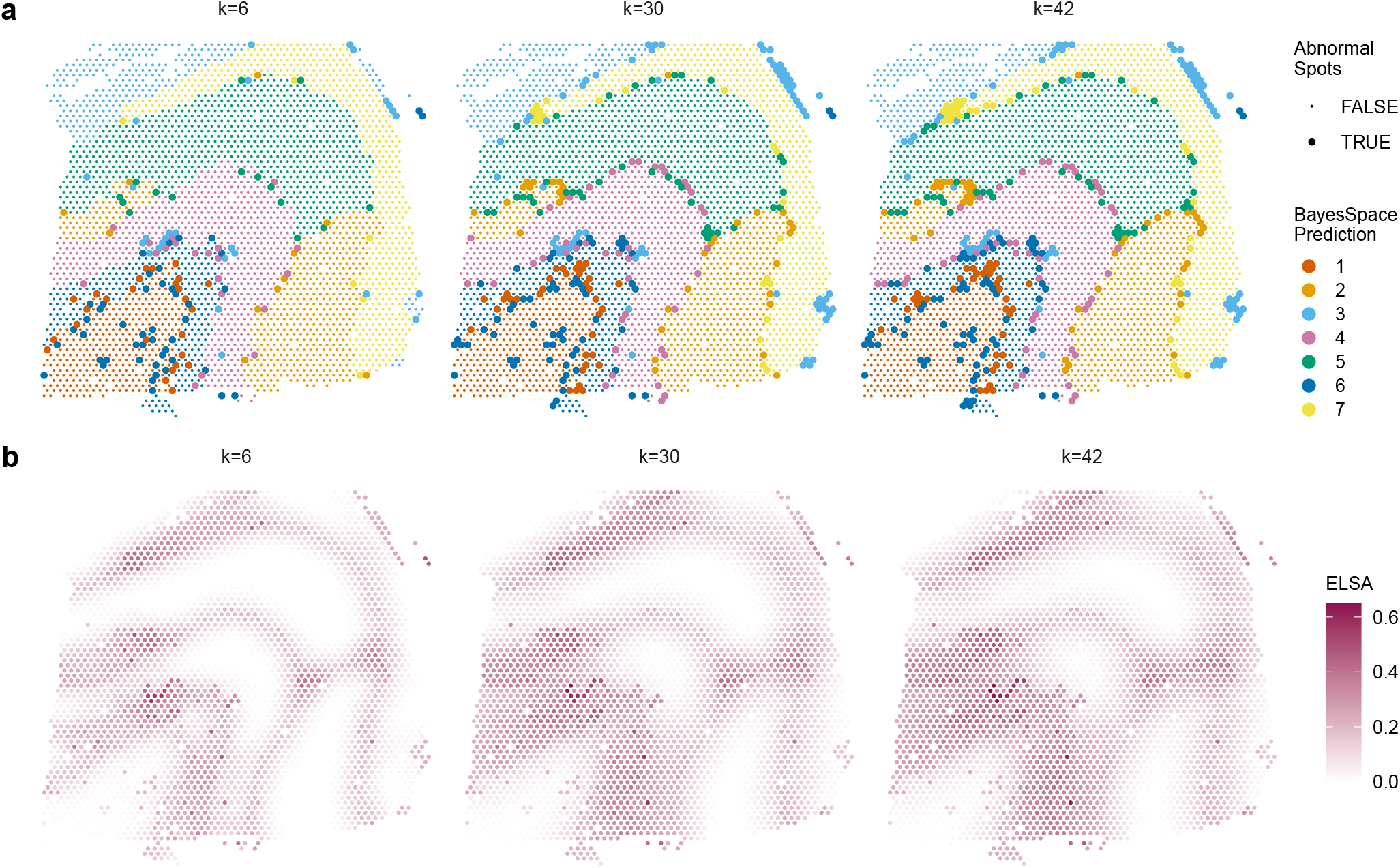
Element-level internal metrics as the size of neighborhood changes. This is related to Figure 7. *k* is the size of the neighborhood used to calculate the element-level metric values. Score in **a** indicates whether a spot is classified as abnormal or not; Score in **b** is the ELSA values.

**Figure S7.**
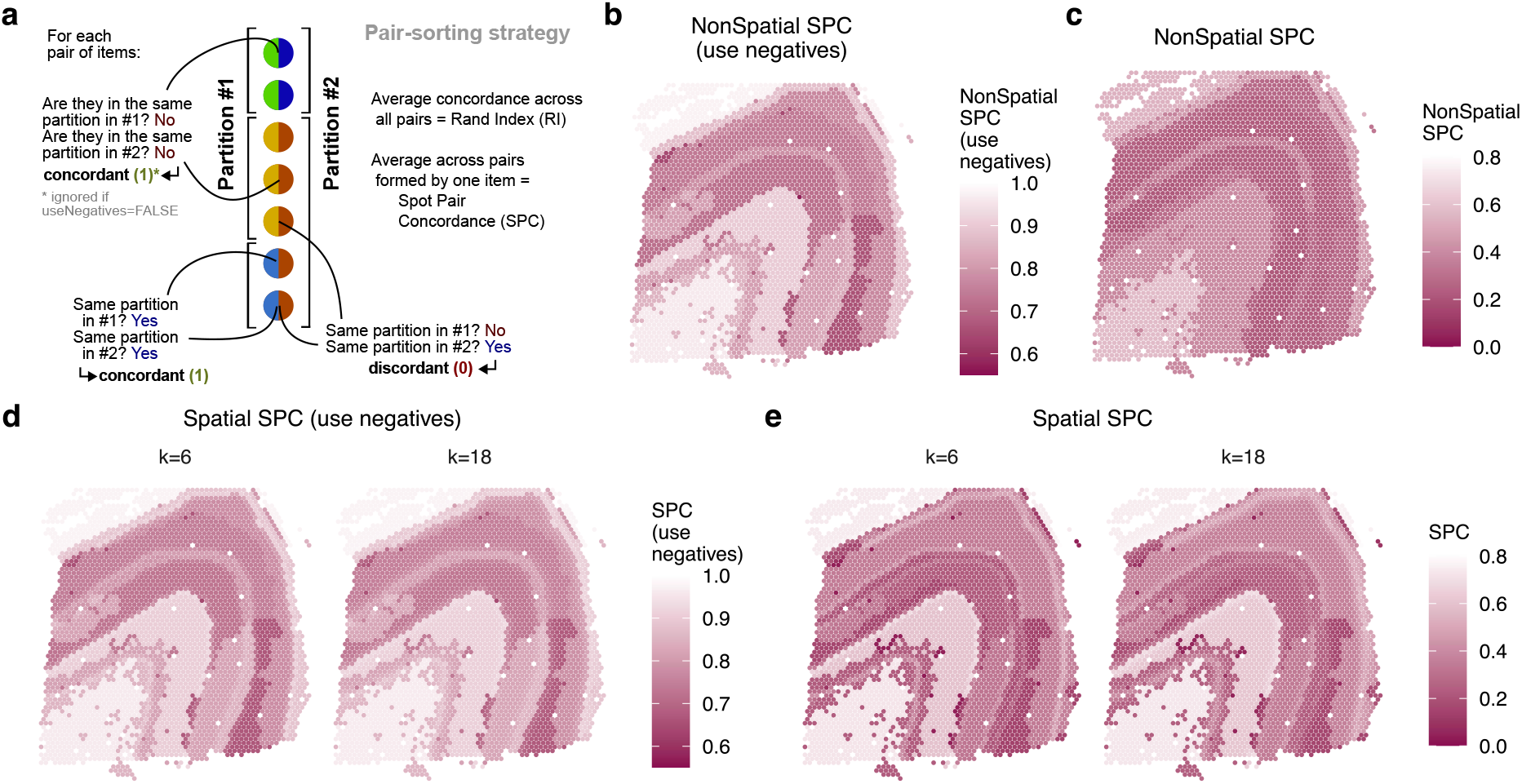
Variants of the Spot-wise Concordance (SPC) score. **a** Illustration of the pair-counting strategy. **b**-**e** show the values of SPC variants for BayesSpace’s prediction as in Figure 7. **b**-**c**: SPC not accounting for spatial relationships, and using **(b)**or not using (**c**) negative pairs. **d**-**e**: Spatially-aware SPC using (**d**) or not using (**e**) negative pairs, and across different neighborhood sizes (*k*).

**Figure S8.**
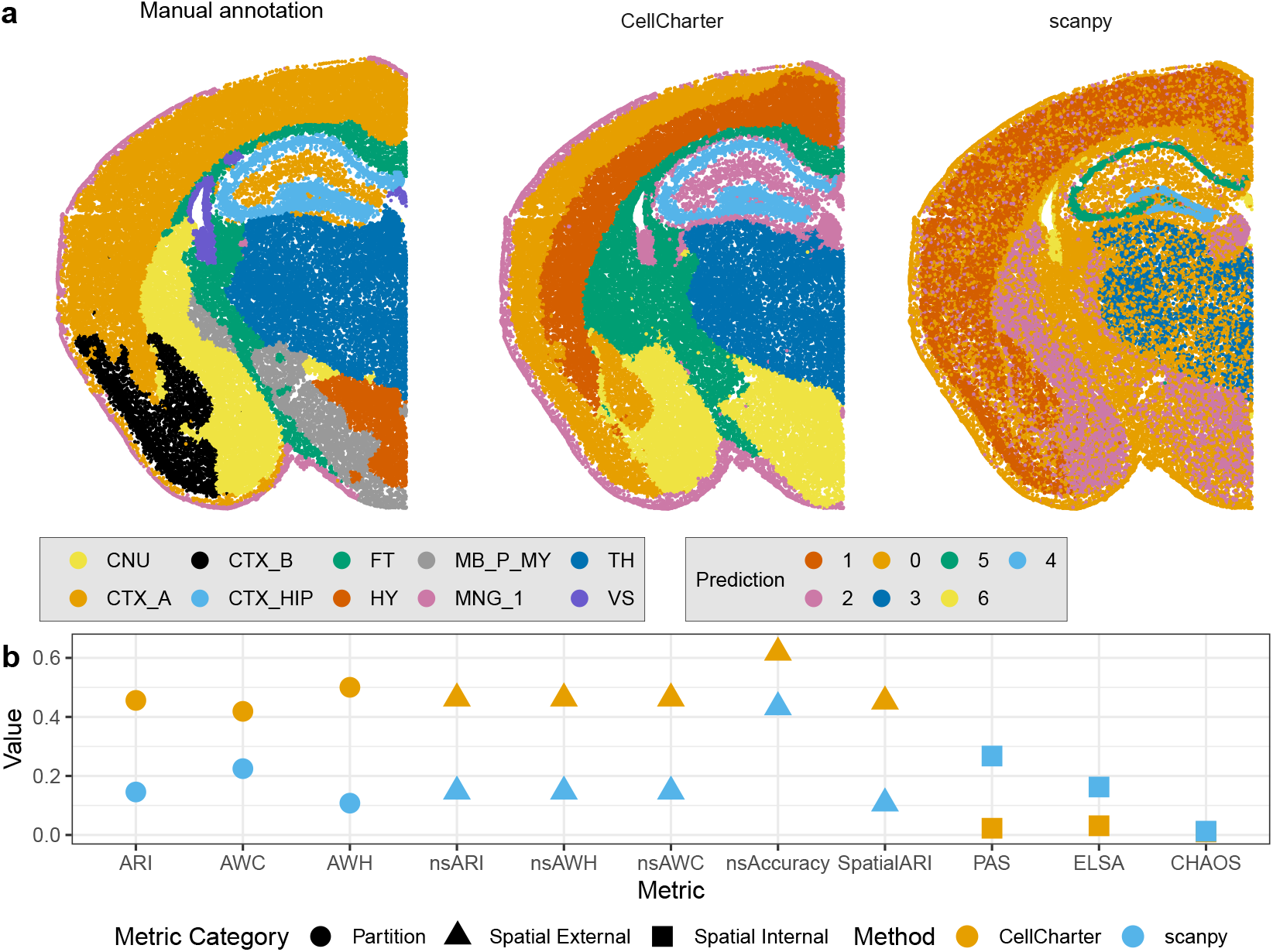
The same metrics are applicable to imaging-based spatial data. To illustrate, the STARmap PLUS mouse brain dataset (slice 11, Shi et al. [2]) is used. In **a**, the first panel shows manual annotations, while the second and third panels show domain predictions from two clustering methods: CellCharter, which is spatially aware, and scanpy, which is not. In **b**, representative partition-based metrics, as well as spatial external and internal metrics are calculated.

**Figure S9.**
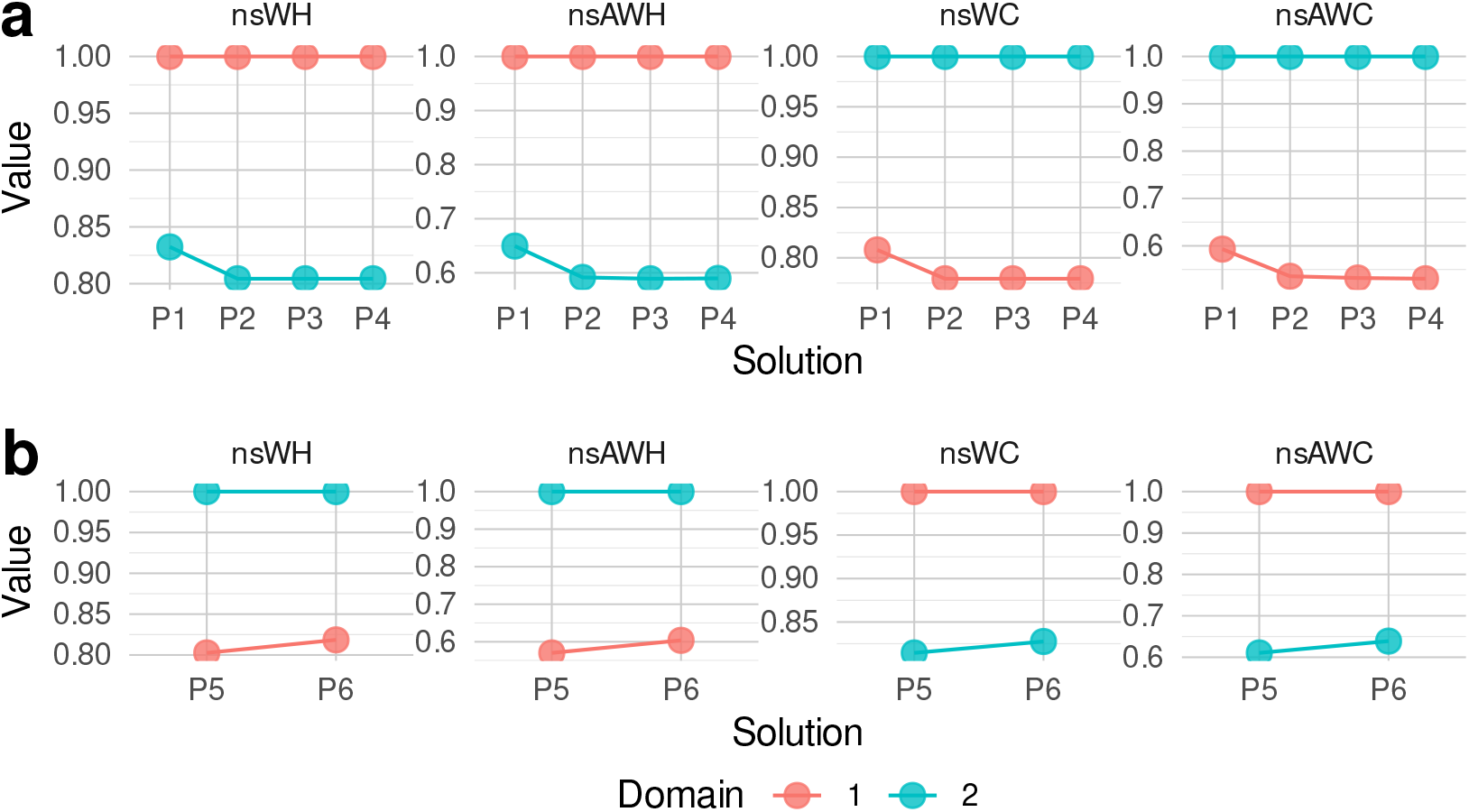
The values of domain-level partition metrics. Related to Figure 8. The cluster-specific (fuzzy-hard version of) spatial WH and AWH, as well as class-specific (fuzzy-hard version of) spatial WC and AWC are calculated for P1-P3 in **a**, and for P4-P5 in **b**, respectively.

**Figure S10.**
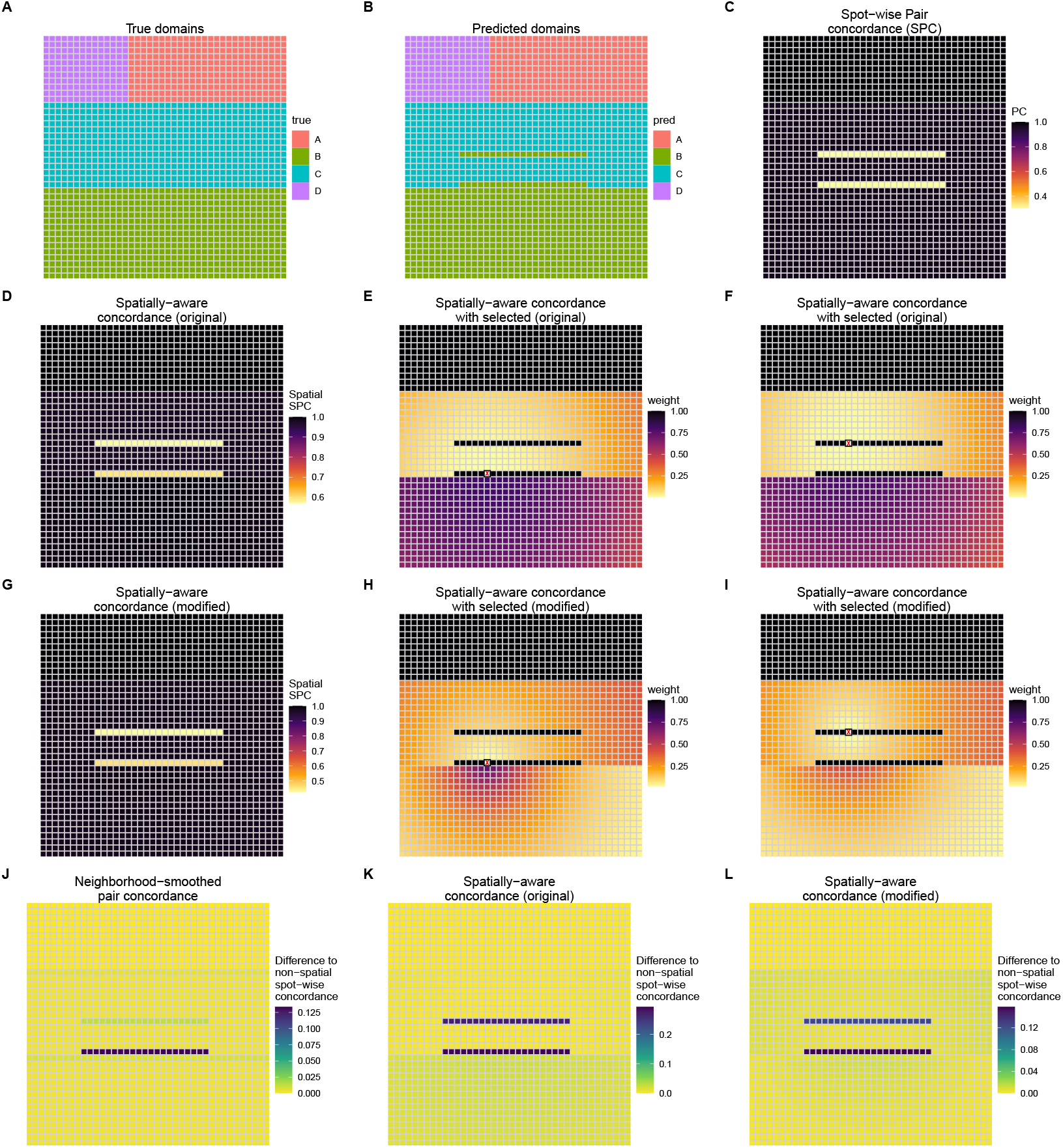
A toy spatial dataset to illustrate the original and adapted distance functions. **a** True domains. **b** Predicted domains. **c** Non-spatial spot-level mean pair concordance (SPC), showing the spots with disagreement between predicted and true domains. **d** Spatially-aware spot concordance, computed using the original distance-weighted functions from Yan et al. [3]. The spot-level differences are very mild, and no difference between the two misclassified stripe can be observed at least by eye. **e-f** Illustration of the distance decay for two different highlighted spots. The colors indicate the (distance-weighed) concordance between each spot and the highlighted spot (marked with an X). While the concordance patterns match the intuition of tolerating the wrong grouping of nearby spots, we believe that this understanding of ‘nearby’ is much to loose, ending up making errors across most of the field of view tolerated. **g-i** Same as **d-f**, but using the adjusted distance functions, which we deem more reasonable. **j-l** The difference between the three spatially-aware spot-wise concordance and the non-spatial one. The neighborhood-smoothed concordance shows good edge tolerance, while the original distance-weighted ARI from Yan et al. [3] does not: the two stripes show a similar increase in concordance from the spatial weighing. In contrast, our modified spatial ARI recovers the difference between the two stripes.

**Figure S11.**
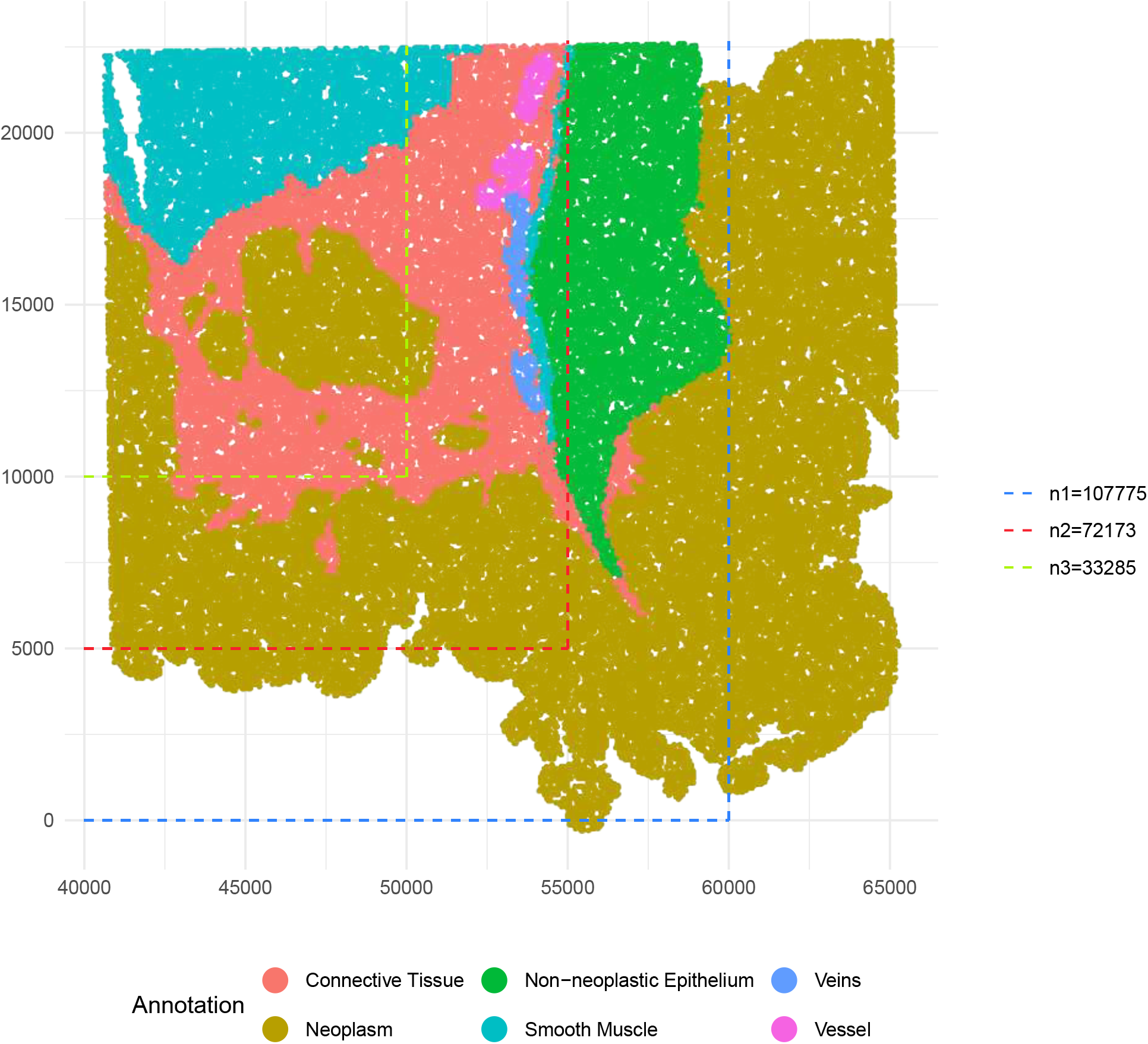
Datasets used for scalability monitoring. The Visium HD colon cancer dataset from [4] was used to monitor CPU time and memory usage in Figure S12. The full slide contains 136,870 cells. To assess scalability, we performed three rounds of subsampling by selecting subregions of the slide, as indicated by the dashed lines, resulting in datasets with 107,775, 72,173, and 33,285 cells, respectively. Each selected metric was then calculated across the three subsampled datasets and the full dataset, repeated five times for each. Cells are colored by the manual annotation from the original paper.

**Figure S12.**
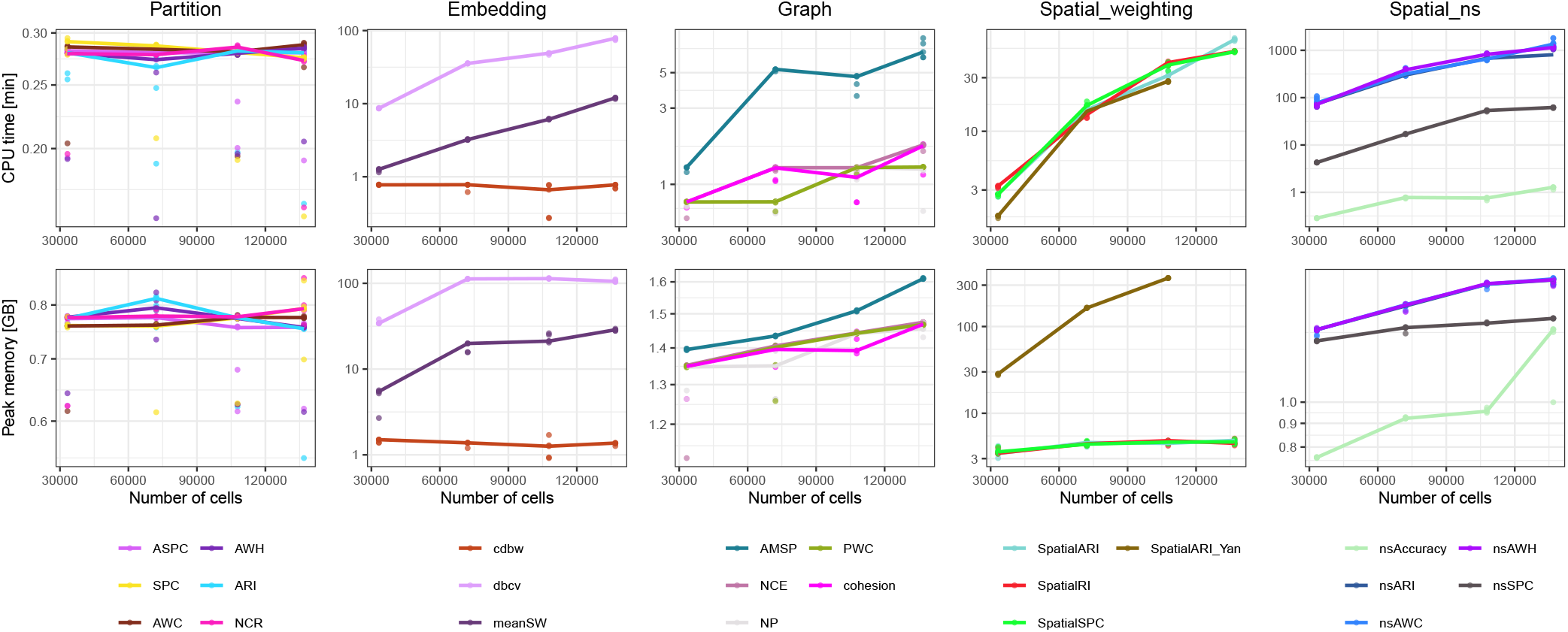
The CPU time and peak memory usage of each selected metric across datasets of different sizes. The Visium HD colon cancer dataset [4] and its subregions (see Figure S11) were used to monitor the runtime and memory usage. Each selected metric was computed on all four dataset sizes, with five repeated runs per size. The individual runs are shown as dots, and the median values across the five runs were used to generate the line plots. The y-axes are presented on a logarithmic scale. The selected metrics are grouped into five categories, each shown in one panel: partition-based, embedding-based, and graph-based metrics, as well as spatial metrics based on distance-weighting (Spatial_weighting), and those based on neighborhood smoothing (Spatial_ns).

## Footnotes

1 In the context of distinguishing populations (as opposed to measuring batch integration), the LISI/SI score of a node is less easy to interpret because it tracks dominance or diversity of the neighborhood independently of whether the neighborhood matches the node’s class.

